# Selective RNA sequestration in biomolecular condensates directs cell fate transitions

**DOI:** 10.1101/2025.05.08.652299

**Authors:** Patrizia Pessina, Mika Nevo, Junchao Shi, Srikanth Kodali, Eduard Casas, Yingzhi Cui, Alicia L. Richards, Emily J. Park, Xi Chen, Florencia Levin-Ferreyra, Erica Stevenson, Nevan J. Krogan, Danielle L. Swaney, Qilong Ying, Qi Chen, Justin Brumbaugh, Bruno Di Stefano

## Abstract

Recent studies have emphasized the significance of biomolecular condensates in modulating gene expression through RNA processing and translational control. However, the functional roles of RNA condensates in cell fate specification remains poorly understood. Here, we profiled the coding and non-coding transcriptome within intact biomolecular condensates, specifically P-bodies, in diverse developmental contexts, spanning multiple vertebrate species. Our analyses revealed the conserved, cell type-specific sequestration of untranslated RNAs encoding key cell fate regulators. Notably, P-body contents did not directly reflect active gene expression profiles for a given cell type, but rather were enriched for translationally repressed transcripts characteristic of the preceding developmental stage. Mechanistically, microRNAs (miRNAs) direct the selective sequestration of RNAs into P-bodies in a context-dependent manner, and perturbing AGO2 or alternative polyadenylation profoundly reshapes P-body RNA content. Building on these mechanistic insights, we demonstrate that modulating P-body assembly or miRNA activity dramatically enhances both activation of a totipotency transcriptional program in naïve pluripotent stem cells as well as the programming of primed human embryonic cells towards the germ cell lineage. Collectively, our findings establish a direct link between biomolecular condensates and cell fate decisions across vertebrate species and provide a novel framework for harnessing condensate biology to expand clinically relevant cell populations.

## MAIN

The ability of stem and progenitor cells to adopt new fates is essential for development and homeostasis in multicellular organisms^8^. Cell fate changes are not primarily driven by alterations in genomic content but instead are orchestrated through regulation at the epigenetic, transcriptomic, and proteomic levels. Post-transcriptional mechanisms are fundamental in regulating gene expression and have emerged as critical factors in establishing and maintaining cell identity^9–11^. For instance, RNA modifications control embryonic and adult stem cell self-renewal by modulating the stability of regulators that dictate cell fate^12–15^. Additionally, RNA splicing, decay, and alternative polyadenylation are pivotal regulatory mechanisms during development, cellular reprogramming, and tissue homeostasis^12, 13, 15–24^.

There is a growing appreciation for the association between RNA regulation and the compartmentalization of transcripts into biomolecular condensates^25–33^. These condensates, which vary in form, function, and location, include P-bodies, evolutionarily conserved structures formed in the cytoplasm through interaction between RNA and RNA-binding proteins (RBPs)^26, 34–41^. Studies in both yeast and mammalian somatic cells demonstrated that untranslated mRNAs can be directed into P-bodies through interaction with proteins involved in translation repression, miRNA-mediated pathways, and mRNA decay processes^35, 39, 42–45^. Elaborating on this point, a recent study in *C. elegans* demonstrated that translationally repressed mRNAs associate to form nanoclusters, which ultimately self-assemble into larger, multiphasic condensates^46^. This study provides a physical model for RNA sequestration into condensates but whether specific RNAs are directed into P-bodies and how this may affect important cellular processes like differentiation remain unknown.

While P-bodies were initially characterized as hotspots for mRNA decay^39, 40^, recent observations have extended our understanding of their regulatory roles, emphasizing their function in sequestering and storing untranslated mRNAs from the translational machinery^47–53^. Supporting this view, repression associated with mRNA accumulation in P-bodies is uncoupled from RNA decay, and P-body transcripts can reenter the ribosome pool upon P-body dissolution^35, 39, 42–45^. For example, the release of translationally repressed mRNAs from P-bodies in *C. elegans* oocytes dramatically increases their concentration in the cytosol (∼9-fold), and their association with translational machinery^46^. These findings suggest that regulation through P-bodies is nuanced and dynamic, providing a precise mechanism to fine tune gene expression.

Emerging evidence suggests a vital role for P-bodies in regulating cell fate. For example, in stem and progenitor cells, P-bodies influence self-renewal by sequestering mRNAs encoding chromatin remodelers and transcription factors^48, 54–56^. Conversely, P-body dysfunction has been implicated in pathologies such as Parkinson’s disease^57^. Aberrant expression of P-body components has also been observed in cancer, suggesting that disruptions to RNA storage and sequestration may destabilize cell identity and contribute to oncogenesis across diverse tissues^40, 58–61^.

Despite these observations, a precise role for RNA sequestration and its associated regulatory mechanisms during differentiation and development remains unclear. Specifically, the extent to which RNAs in P-bodies are conserved across various cell identities and species, as well as the molecular mechanisms governing their sequestration, remains poorly understood. Furthermore, investigating whether acute perturbations in P-body assembly affect cell identity could provide key insights into the role of RNA condensates in cell fate determination and their potential applications in generating clinically relevant cell types.

To address this important gap in knowledge, we purified intact P-bodies and quantified constituent coding and non-coding RNAs during differentiation across all three primary germ layers, during reprogramming to pluripotency, as well as in terminally differentiated cells. These conditions encompass multiple developmental contexts and vertebrate species. Our findings revealed that RNA sequestration in P-bodies is a conserved regulatory mechanism, leading to the storage of specific transcripts that reflect preceding developmental stages. Using a degron system, we demonstrated that the ribosome occupancy of mRNAs enriched in P-bodies, including fate-instructive transcripts, increased following acute dissolution of P-bodies. Large-scale quantitative proteomics confirmed increased translation for these transcripts. Mechanistically, we demonstrated a critical role for miRNAs in regulating RNA sequestration in a context-dependent manner. Building on these insights, we showed that perturbing RNA sequestration can both activate a totipotency transcriptional program in naïve human pluripotent cells and facilitate the efficient conversion of primed human embryonic stem (ES) cells in primordial germ cell-like cells. Collectively, our findings define a fundamental and conserved role for P-bodies in cell fate specification and highlight novel strategies to exploit RNA condensates for directing cell identity.

## RESULTS

### Purification of intact P-bodies

To profile the content of RNA condensates, we adapted a fluorescence-activated sorting method for purifying intact P-bodies^49, 62^ (**Fig. 1a**). As proof of concept, we transduced HEK293T cells with lentivirus expressing either a GFP-LSM14A construct or a GFP control. LSM14A is an established protein component of P-bodies^49^ and GFP-LSM14A puncta colocalized with the P-body marker EDC4^63^, confirming the specificity of our construct (**Fig. 1b**). Following cell lysis, intact GFP-LSM14A particles were detectable by epifluorescence imaging and were readily isolated through Fluorescence-Activated Particle Sorting (FAPS), exclusively from GFP-LSM14A-expressing cells (**Fig. 1c and Extended Data Fig. 1a**). By contrast, cells expressing the cytoplasmic GFP control construct had no detectable particles for sorting (**Fig. 1c and Extended Data Fig. 1a**).

**Figure 1.**
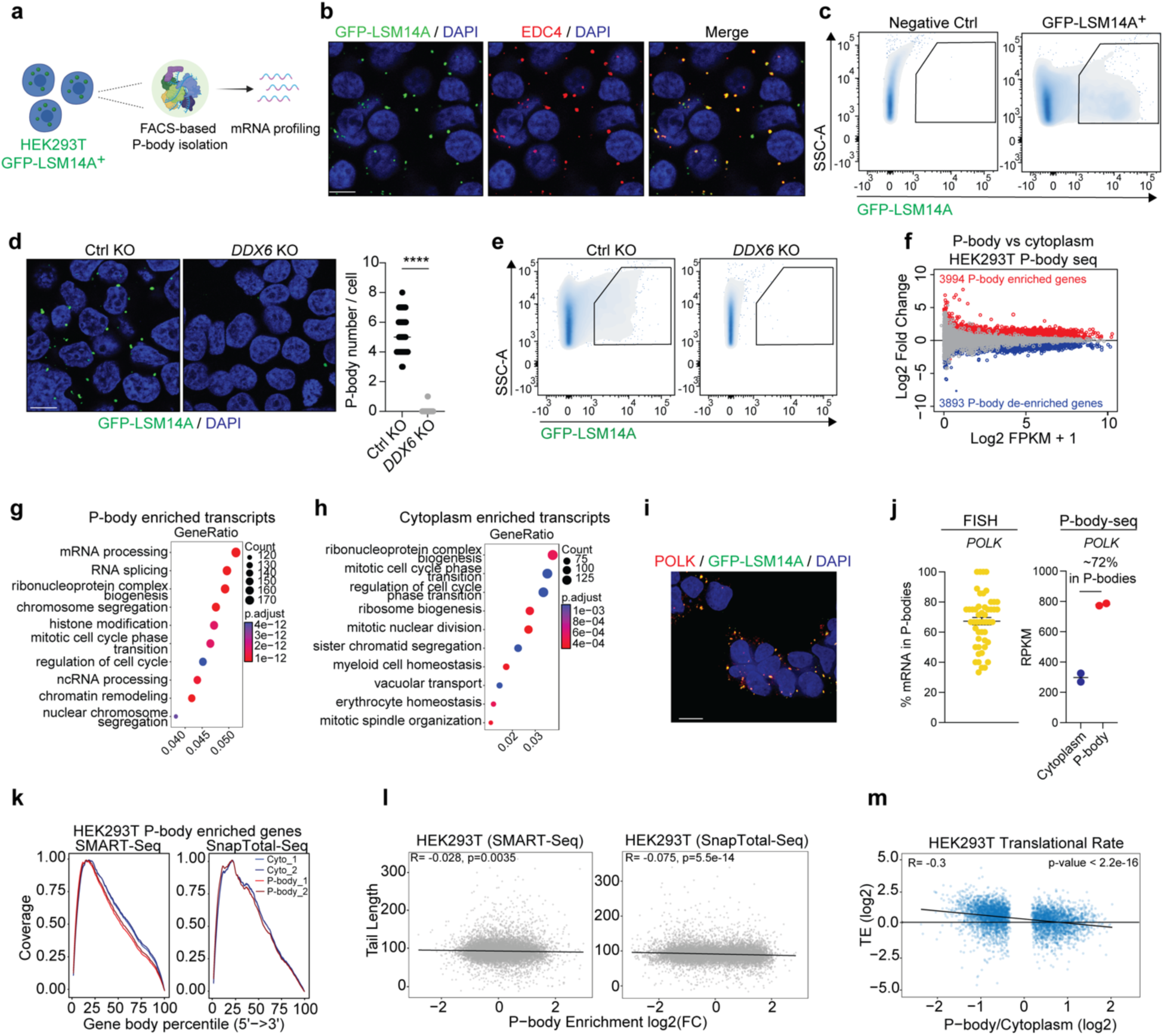
P-body-seq permits comprehensive profiling of P-body contents. **(a)** Schematic for the purification and transcriptomic profiling of P-bodies from HEK293T cells based on the expression of GFP-LSM14A. **(b)** Representative IF imaging of GFP-LSM14A puncta (green), colocalizing with EDC4 puncta (red) in HEK293T cells. Nuclei were counterstained with DAPI (blue) (scale: 10mm). **(c)** Representative flow cytometry plots showing gating for GFP-LSM14A+ P-bodies in HEK293T cells. **(d)** Representative imaging of GFP-LSM14A puncta (green) in control and *DDX6* KO HEK293T cells. Nuclei were counterstained with DAPI (blue) (scale: 10mm) (left panel). P-body number in control (n=50 cells) and *DDX6* KO (n=50 cells) HEK293T cells (right panel). Unpaired Student’s t-test, mean ± s.d., ****: p<0.0001. **(e)** Representative flow cytometry plots showing gating for GFP-LSM14A+ P-bodies in control and *DDX6* KO HEK293T cells. **(f)** MA plot of RNA-seq data depicting P-body enriched genes in red and cytoplasm enriched genes in blue in HEK293T cells (n=2, p < 0.05). **(g)** GO pathway analysis of P-body enriched mRNAs in HEK293T cells. **(h)** GO pathway analysis of cytoplasmic fraction-enriched mRNAs in HEK293T cells. **(i)** Representative FISH imaging of *POLK* RNA molecules (red) combined with imaging of GFP-LSM14A puncta (green). Nuclei were counterstained with DAPI (blue) (scale: 10mm). **(j)** Quantification of *POLK* mRNA molecules in P-bodies based on FISH (n=50 cells; right) and P-body sequencing (right). **(k)** Read coverage distribution over the gene body of the longest annotated isoforms for genes enriched in P-bodies or cytoplasm in HEK293T cells. **(l)** PolyA tail length as determined in4 compared to P-body enrichment based on SMART-Seq and SnapTotal-Seq, Pearson correlation test. **(m)** Translation efficiency (log2 (Ribo-seq counts/RNA-seq counts)) negatively correlates with mRNA enrichment in P-bodies in HEK293T cells, Pearson correlation test.

To further corroborate FAPS-based purification of P-bodies, we depleted *DDX6* in GFP-LSM14A HEK293T cells to disrupt P-body assembly and maintenance^40, 48, 52, 64^. Both shRNA-mediated suppression of *DDX6* and CRISPR knockout resulted in the complete loss of GFP^+^ particles based on immunofluorescence and flow cytometric analyses, again suggesting that GFP-LSM14A specifically permits the identification and isolation of P-bodies (**Fig. 1d, e and Extended Data Fig. 1b, c**). Finally, to confirm the presence of RNA in GFP-LSM14A particles, we stained lysates from GFP-LSM14A expressing cells with SYTOX BLUE, a fluorescent nucleic acid dye capable of binding RNA. We observed RNA labeling in GFP^+^ particles, while SYTOX BLUE signal was negligible in GFP^-^ particles (**Extended Data Fig. 1d**). Together, these findings confirm the robust labeling of P-bodies by GFP-LSM14A, which can be further isolated intact via FAPS.

We next characterized the RNA content of the purified P-bodies and corresponding cytosolic fractions using SMART-seq^65^ (hereafter defined as P-body-seq). In total, we detected 3994 mRNAs preferentially associated with P-bodies in HEK293T cells relative to the cytosol (**Fig. 1f**). Gene Ontology (GO) analysis revealed that P-body-enriched mRNAs encoded pivotal regulators of RNA processing, transcription, chromatin organization, and the cell cycle (**Fig. 1g**), while cytosolic mRNAs were predominantly involved in housekeeping functions such as ribosome assembly and organelle structural components (**Fig. 1h**). Thus, P-bodies appear to sequester RNAs encoding key chromatin regulatory factors and proteins associated with RNA processing. To ensure that our analyses were not biased by polyA-directed library preparations, we performed P-body-seq using SnapTotal-Seq^66^, an alternative low-input library preparation method that utilizes random primers rather than oligo(dT). SnapTotal-Seq results were highly consistent with SMART-Seq data, with approximately 70% overlap in P-body-enriched transcripts (**Extended Data Fig. 1e**). Moreover, we compared our dataset to P-body enriched transcripts previously reported in HEK293T cells^49^, and found that 3,038 mRNAs (76%) are shared between datasets (**Extended Data Fig. 1f**). To further confirm our P-body-seq results using an orthogonal approach, we employed single-molecule fluorescence *in situ* hybridization (smFISH) in conjunction with immunofluorescence to visualize the subcellular localization of individual RNA molecules^67^. Focusing on *POLK*, one of the transcripts enriched in our P-body-seq analysis, we confirmed its localization within P-bodies (**Fig. 1i, j**). In line with previous reports^46, 58^, approximately 70% of total *POLK* mRNA was localized to P-bodies using either smFISH or P-body-seq, demonstrating that substantial and functionally relevant levels of transcripts may be localized to P-bodies.

Since P-bodies have been proposed as sites for RNA decay^39, 40, 68, 69^, we assessed whether transcripts in HEK293T cells were targeted to P-bodies for degradation. Analysis of read distributions revealed no evidence of truncated transcripts in P-bodies (**Fig. 1k**), in line with previous reports^46, 49, 58^. In addition, we observed poor correlation between mean poly(A) tail length^4^ and P-body enrichment (r=-0.028 in SMART-Seq data and r=-0.075 in SnapTotal-Seq data) (**Fig. 1l**), indicating that P-body-localized RNAs are intact and do not undergo preferential deadenylation relative to cytoplasmic transcripts.

We next asked whether properties intrinsic to certain RNAs correlated with their enrichment in P-bodies. While transcript length had no appreciable relationship with P-body localization, we found that transcripts with high AU content were enriched amongst P-body-targeted mRNAs (**Extended Data Fig. 1g**). Notably, AU-rich sequences are associated with inefficient translation^48, 70, 71^. Accordingly, analysis of previously published ribosome profiling data obtained in HEK293T cells^4^ revealed reduced translation efficiency for P-body-associated transcripts (**Fig. 1m**). These data suggest that transcripts sequestered in P-bodies are translationally repressed compared to cytoplasmic-enriched mRNAs. Collectively, these findings validate our methodology for isolating intact P-bodies and reveal the sequestration of untranslated mRNAs encoding pivotal RNA and chromatin binding proteins within these subcellular condensates in mammalian cells.

### P-body contents are cell type-specific

Transcripts localized to P-bodies can re-enter translation and contribute to protein-level expression in response to genetic manipulation or environmental stimuli^44, 49, 64, 70, 72, 73^. This observation raises the intriguing possibility that the sequestration or release of fate-instructive transcripts is a fundamental regulator of cell fate decisions during development. However, establishing a functional role for P-bodies in regulating complex biological processes (e.g., development) is challenging, in part because comprehensive, sequencing-based approaches to identify the contents of P-bodies have thus far been limited to non-vertebrates or transformed cell lines^44, 49, 74^. To address this need, we applied our approach to profile P-body-enriched transcripts to a panel of human cell types across a range of developmental stages. We engineered human ES cells to express a GFP-LSM14A knock-in construct from the *AAVS1* locus and cultured them under naïve and primed conditions. These cells represent the pre- and post-implantation epiblast, respectively^75, 76^. We also induced lineage-specific differentiation of the same human pluripotent stem cell line into progenitors of all three germ layers (mesoderm and endoderm progenitors, as well as neural progenitors), representing multipotent states, and ultimately into neurons (**Fig. 2a**). Immunofluorescence based on markers for each cell type confirmed robust differentiation into the expected lineages (80% efficiency or greater; (**Extended Data Fig. 2a**)), which was supported by the expression of a broader set of lineage-specific genes (**Extended Data Fig. 2b**). We detected EDC4^+^ P-bodies in each cell type, albeit in varying amounts (**Extended Data Fig. 2c, d**), suggesting differences in RNA sequestration between cell types.

**Figure 2.**
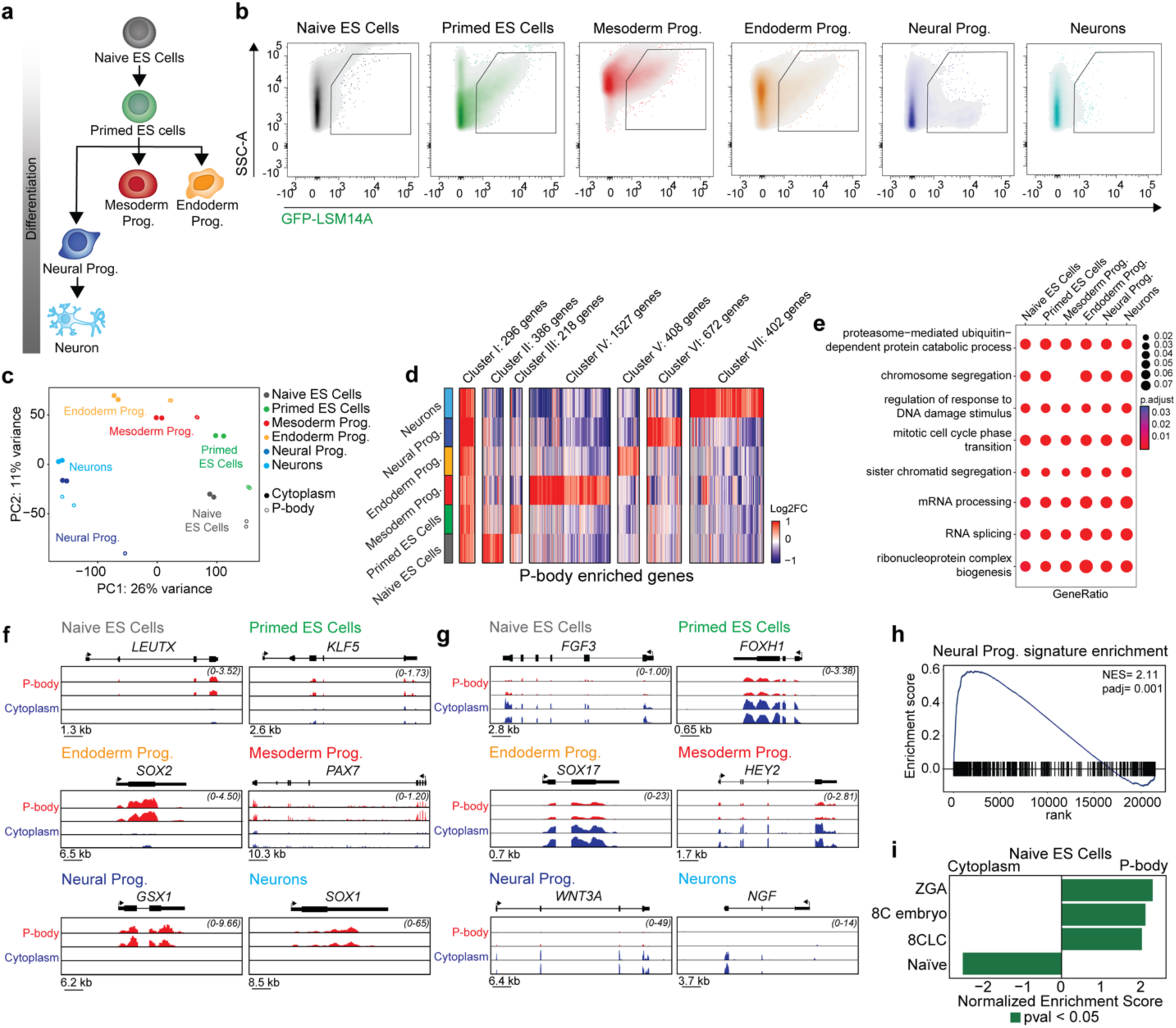
RNA sequestration in P-bodies is cell-type specific. **(a)** A schematic highlighting developmental stages profiled by P-body-seq. **(b)** Representative flow cytometry plots showing gating for GFP-LSM14A+ P-bodies in the indicated samples. **(c)** Principal component analysis of RNA-seq data for the indicated samples. **(d)** Heatmap showing expression levels of differentially enriched mRNAs between purified P-body fractions of the indicated samples. Gene number in each cluster is indicated in the Fig. (n=2, p < 0.05). **(e)** GO pathway analysis of P-body enriched mRNAs in the indicated samples. **(f, g)** Gene tracks showing individual genes from RNA-seq data. **(h)** GSEA analysis for a neural progenitor-related gene expression signature in purified P-bodies from human neurons. **(i)** Normalized Enrichment Score (NES) of genes in gene sets from2, 3, p<0.05.

To identify P-body contents, we sorted GFP-LSM14A^+^ particles from each cell type, isolated RNA, and performed SMART-seq (**Fig. 2b**). Using this approach, we identified a broad range of RNA biotypes (protein coding mRNAs, lincRNAs, etc.), confirming their presence in P-bodies (**Extended Data Fig. 2e**). We note, however, that RNA sequestration in P-bodies is not merely a reflection of overall expression level, as P-body enriched transcripts were distributed similarly amongst expression quartiles (**Extended Data Fig. 2f**). We likewise identified repetitive elements in P-bodies, though we did not detect marked differences in their abundance between P-bodies and cytoplasm (**Extended Data Fig. 2g, h**). Based on principal component analysis (PCA), we observed distinct mRNA profiles for P-bodies from each cell type, suggesting that distinct cell identities have a unique set of transcripts stored in P-bodies (**Fig. 2c**). PCA further suggested a stepwise progression from naïve ES cells towards mesoderm/endoderm lineages on one path and the ectoderm lineage on another (**Fig. 2c**). Surprisingly, we found that P-body contents did not necessarily cluster most closely with the cytoplasm from the same cell type (**Fig. 2c**). Instead, P-body-enriched transcripts of differentiated progeny often clustered most closely with the cytoplasmic transcripts of stem or progenitor cells from the preceding differentiation state. For example, P-body samples from neurons cluster most closely with cytoplasmic samples from neural progenitors and P-bodies contents of mesoderm progenitors cluster together with the cytoplasm of primed ES cells (**Fig. 2c**). These data suggest that perhaps P-bodies sequester transcripts from stem and progenitor cells to suppress their protein-level expression during differentiation.

### P-body contents reflect preceding human developmental stages

We next grouped transcripts based on their enrichment across our panel of cell types (**Fig. 2d and Extended Data Fig. 2i**). A small fraction of transcripts was enriched in the P-bodies of all cell types and functional annotation revealed that this gene set was enriched for gene ontology categories linked to DNA damage and cell cycle progression (**Fig. 2e**). By contrast, transcripts enriched in the P-bodies of single cell types encoded fate-instructive factors, including lineage-specific transcription and chromatin factors, such as *SOX1*^77^ for neurons, *LEUTX*^78^ in naïve ES cells, and *KLF5*^79^ in primed ES cells (**Fig. 2f**). In line with PCA, these factors were generally associated with the preceding developmental stage. For instance, SOX1 is a transcription factor that plays a role in the self-renewal of neural progenitor cells and is downregulated during neuronal specification^80, 81^. Correspondingly, *SOX1* transcripts were heavily enriched in neuronal P-bodies (**Fig. 2f**). On the other hand, transcripts enriched in the cytoplasm directly reflected the identity and function of the current cell state (e.g., *SOX17* in endoderm progenitors; **Fig. 2g**). These data further support the notion that cell type-associated transcripts remain available in the cytoplasm, while transcripts characteristic of preceding developmental stages are sequestered into P-bodies.

To extend these observations, we conducted gene set enrichment analysis (GSEA) on P-body-associated RNAs in neurons, which revealed the robust enrichment of a transcriptional signature characteristic of neural progenitor cells (**Fig. 2h**). In line with these data, P-body dissolution impaired neuronal specification from neural progenitors^48, 54, 56^, providing functional evidence that P-bodies sequester genes related to stem and progenitor cell self-renewal to facilitate cell fate changes. To investigate whether our observations in the ectoderm lineage extend to mesoderm and endoderm, we performed GSEA of P-body-associated RNAs in mesoderm and endoderm progenitors. This analysis revealed consistent enrichment of transcripts from the preceding, primed ES cell state in both lineages (**Extended Data Fig. 3a**). To further assess the functional requirement of P-bodies in endoderm differentiation, we employed a CRISPRi-based approach to deplete DDX6^48^, an essential P-body component^52, 64, 82^, which induced dissolution of P-bodies in endoderm progenitors (**Extended Data Fig. 3b**). Notably, DDX6 depletion significantly impaired endoderm progenitor differentiation toward AFP^+^ hepatocytes (**Extended Data Fig. 3c**). Moreover, DDX6-depleted endoderm progenitor cells failed to induce other hepatic differentiation markers such as *ALBUMIN*, *G6PC* and *CYP3A7* compared to control cells (**Extended Data Fig. 3d**). These results suggest that P-bodies are essential for differentiation through the sequestration of transcripts from the preceding developmental state.

We next reasoned that if P-body-mediated sequestration of stem and progenitor cell transcripts reinforces the early phases of differentiation, these transcripts would eventually decline in P-bodies as differentiation completes and transcriptional profiles shift to lock in the differentiated state. To test this possibility, we extended the culture of neurons in our differentiation system to 20 days (**Extended Data Fig. 3e**). Analysis of P-body content in mature neurons showed substantial overlap (∼50%) with transcripts sequestered from neurons at day 7 of differentiation (**Extended Data Fig. 3f**); however, prolonged culture led to a progressive depletion of neural progenitor-associated transcripts from P-bodies (**Extended Data Fig. 3g**). These data indicate that RNAs in P-bodies of post-mitotic neurons gradually decrease as terminal differentiation is achieved. To test whether this observation was consistent in an alternate lineage, we generated mature endoderm progenitors (**Extended Data Fig. 3h**). Compared to endoderm progenitors generated directly from primed ES cells, P-bodies from mature endoderm progenitors demonstrated decreased enrichment for a gene expression profile reflective of the preceding developmental stage (i.e., primed ES cells; **Extended Data Fig. 3i)**. This progression suggests that P-bodies may serve as temporary reservoirs for developmental transcripts, potentially maintaining cellular plasticity during early development—a function that becomes dispensable as cells achieve terminal differentiation.

We next examined P-body-associated transcripts across two pluripotent states. In primed ES cells, P-bodies showed enrichment of naïve ES cell-specific gene signatures (**Extended Data Fig. 3j**), consistent with previous observations that P-body dissolution promotes primed-to-naïve state conversion^48^. Similarly, P-bodies in naïve ES cells were enriched for gene expression signatures characteristic of the preceding developmental stage, including transcripts associated with zygotic genome activation (ZGA), the 8-cell embryo, and 8C-like cells (8CLC)^3, 83^ (**Fig. 2i and Extended Data Fig. 3k**). While naïve human ES cells typically express low levels of ZGA- and 8CLC-related transcripts^3^, our data suggest these transcripts are sequestered into P-bodies to prevent their translation and inhibit inappropriate reversion to an earlier developmental state. By contrast, a gene expression signature related to naïve pluripotency was heavily enriched in the cytoplasm relative to P-bodies (**Fig. 2i**), likely reflecting active expression of genes that support the naïve cell fate. These findings suggest that RNA condensates in ES cells play a critical role in selectively sequestering key totipotency-associated factors. Altogether, these data indicate that P-body contents do not simply reflect the gene expression profiles of a given cell type but are instead enriched for transcripts characteristic of the preceding developmental stage.

### RNA sequestration is conserved across vertebrates

Our discovery that P-bodies play a fundamental role in sequestering RNAs encoding fate-instructive proteins across human cell types prompted us to investigate whether P-body contents and functions are conserved across vertebrate species. To this end, we first considered murine naïve and primed ES cells^84, 85^. Similar to their human counterparts, naïve mouse ES cells resemble the *in vivo* pre-implantation mouse epiblast, while the primed state represents post-implantation epiblast cells^86^. Conversion from naïve to primed mouse ES cells was efficient, as evidenced by distinct stage-specific morphological and transcriptional changes^85^ (**Fig. 3a, b and Extended Data Fig. 4a**). Following transduction with a GFP-LSM14A lentivirus, both naïve and primed ES cells harbored GFP-LSM14A^+^ P-bodies in similar numbers (**Fig. 3c**).

**Figure 3.**
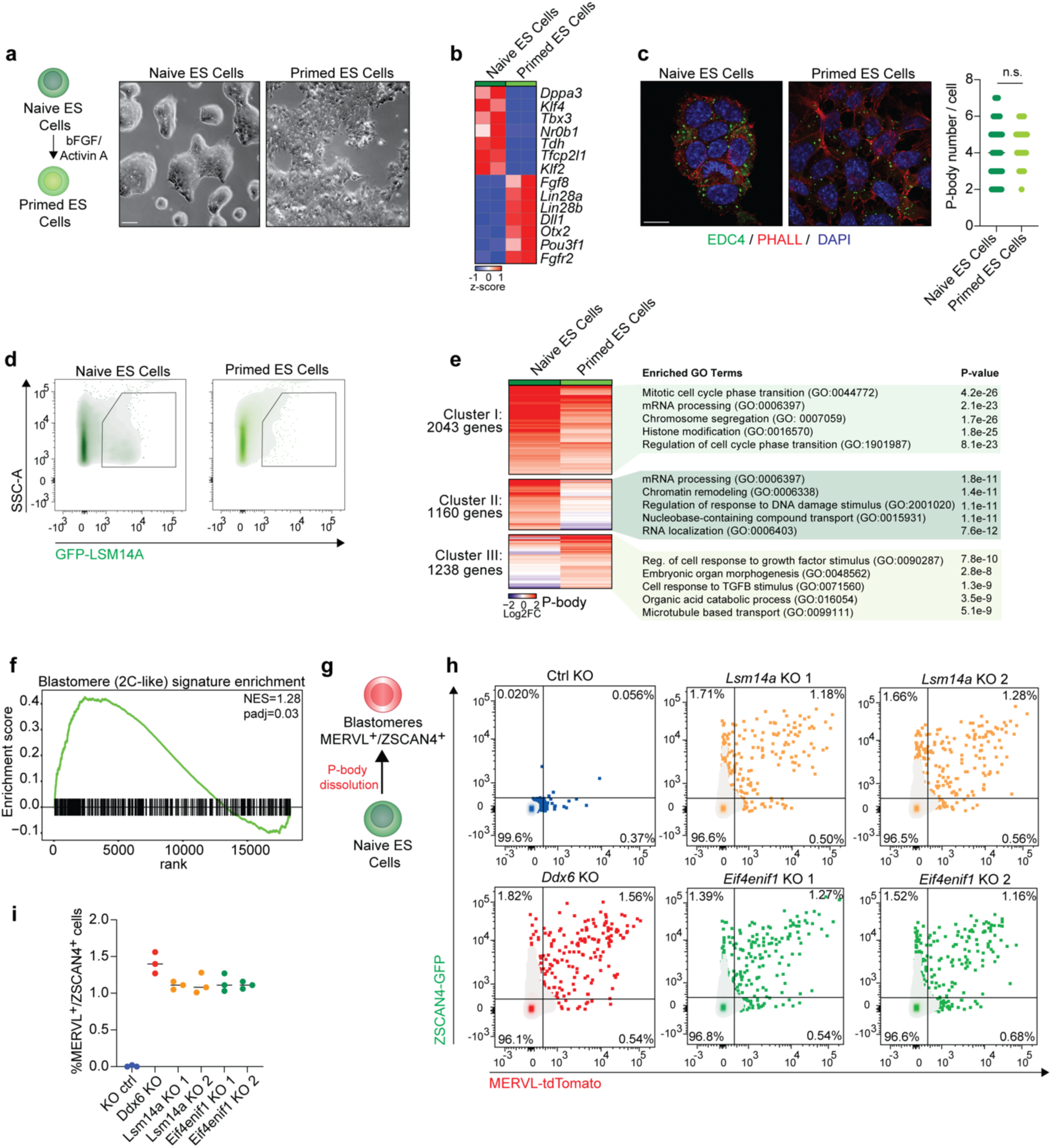
RNA sequestration in P-bodies safeguards mouse ES cell identity. **(a)** Schematic of the conversion from naïve to primed state mouse ES cells (left panel). Representative bright field images of naïve and primed mouse ES cells (scale: 50μm) (right panel). **(b)** Heatmap showing expression levels of naïve-specific and primed-specific transcripts. (n=2). **(c)** Representative IF imaging of EDC4 puncta (green) in naïve and primed mouse ES cells. Cell membranes were labeled with Phalloidin (red) and nuclei were counterstained with DAPI (blue) (scale: 10μm) (left panel). P-body number in naïve (n=60 cells) and primed (n=60 cells) mouse ES cells (right panel). Unpaired Student’s t-test, mean ± s.d., n.s.: p >0.05. **(d)** Representative flow cytometry plots showing gating for GFP-LSM14A+ P-bodies in naïve and primed mouse ES cells. **(e)** Heatmap showing expression levels of differentially enriched mRNAs between purified P-body fractions of naïve and primed mouse ES cells, with GO pathway analysis of P-body-enriched transcripts for the indicated clusters. Gene number in each transcript cluster is indicated in the figure (n=2, p < 0.05). **(f)** GSEA analysis of blastomere-related genes in purified P-body fraction vs. cytoplasmic fraction from naïve mouse ES cells, NES=1.28, p=0.006. **(g)** A schematic of the strategy for P-body dissolution in mouse naïve ES cells carrying blastomere-specific reporters (MERVL-tdTomato and ZSCAN4-GFP). **(h, i)** Flow cytometric quantification of MERVL-tdTomato+ and ZSCAN4-GFP+ cells upon *Lsm14a*, *Ddx6,* and *Eif4enif1* KO in mouse naïve ES cells. Control (n=3), *Lsm14a* (n=3), *Ddx6 KO* (n=3) and *Eif4enif1* KO (n=3).

To compare the P-body transcriptome in mouse naïve and primed ES cells, we conducted P-body purification (**Fig. 3d**) followed by SMART-seq analysis of both P-body and cytoplasmic fractions. Extending and reinforcing our results in human pluripotent stem cells, our analysis revealed that, while a subset of RNAs was commonly enriched in P-bodies in both cell states, a significant fraction of P-body-enriched transcripts exhibited cell-type-specific localization (54%) (**Fig. 3e and Extended Data Fig. 4b**). These results confirm that RNA sequestration is orchestrated in a cell-type-specific manner in mice, potentially contributing to cell fate transitions. To explore this possibility further, we performed GO analysis and found that transcripts enriched in P-bodies of both naïve and primed ES cells were associated with cell cycle progression and RNA processing, similar to human datasets (**Fig. 3e**). Likewise, transcripts that sequestered exclusively in P-bodies of naïve and primed ES cells encoded proteins with functions related to chromatin remodeling and growth factor response, respectively (**Fig. 3e**). Together, these data further support the idea that P-bodies in different cell types across species sequester distinct subsets of transcripts related to relevant biological processes.

To directly assess cross-species conservation, we then compared the mRNA orthologs sequestered within P-bodies of pluripotent cells between mouse and human in both the naïve and primed states. Remarkably, we found that approximately 50% of P-body-enriched transcripts are shared between mouse and human naïve ES cells as well as primed ES cells (**Extended Data Fig. 4c, d**). This observation led us to ask whether this mechanism is also conserved across phylogenetically distant vertebrate species. To test this possibility, we transduced chicken ES cells with our GFP-LSM14A lentivirus. Chicken ES cells are derived at the blastoderm stage and can contribute to chimeric embryos^87^. In line with our findings in human and mouse cells, we detected GFP-LSM14A particles in chicken ES cells (**Extended Data Fig. 4e**), which we purified by FAPS. Following P-body-seq, we found that mRNAs enriched in P-bodies of chicken ES cells exhibit extensive overlap with the P-body transcriptomes of mouse and human (**Extended Data Fig. 4f**). Comparative analysis further revealed similarities in transcript features among chicken, mouse, and human ES cells, particularly in the preferential sequestration of AU-rich mRNAs in P-bodies, regardless of transcript length (**Extended Data Fig. 4g**). Thus, RNA sequestration in P-bodies appears to be conserved across vertebrates.

To further analyze the conservation of RNA sequestration across vertebrates, we assessed the strength of association among P-body transcripts from all species. Utilizing the Odds Ratio statistic, we identified a significant positive association of chicken ES cells with both mouse and human ES cells, with the strongest association observed between mouse and human samples (**Extended Data Fig. 4h**). Furthermore, GO analysis revealed enrichment in categories related to RNA processing, transcription, chromatin organization, and cell cycle among P-body-associated transcripts that are shared between chicken ES cells and mouse/human naïve and primed ES cells (**Extended Data Fig. 4i**). These data corroborate and extend our findings in human and mouse ES cells, indicating that transcripts sequestered in P-bodies of chicken ES cells encode important regulatory factors. Collectively, our data indicate that P-body-enriched transcripts are distinct between cell types and underscore the conservation of RNA sequestration as a regulatory mechanism across vertebrates.

### Loss of RNA sequestration in P-bodies induces translation of proteins related to totipotency in murine ES cells

In our analysis of human pluripotent stem cells, we found that P-bodies sequester transcripts related to the 8-cell stage embryo and zygotic genome activation (**Fig. 2i**). In mice, the equivalent developmental stage occurs during the 2-cell embryo, although rare cells in naïve mouse ES cell cultures, known as 2C cells, transiently express genes characteristic of the 2-cell state^5^. To determine whether transcripts enriched in the P-bodies of mouse naïve ES cells comprise a signature characteristic of totipotent cells, we performed GSEA. Consistent with our findings in human naïve ES cells, we observed a significant enrichment of a 2C-associated gene expression signature among P-body-enriched mRNAs, regardless of library preparation method (**Fig. 3f and Extended Data Fig. 5a**). This observation further indicates that P-body-based regulation of developmental processes is conserved between species. To test the functional consequence of sequestration of 2C transcripts, we employed CRISPR-Cas9 to knockout

*Ddx6* and dissolve P-bodies in mouse ES cells harboring 2C-specific reporters, *Mervl-tdTomato* and *Zscan4-GFP*^88, 89^ (see schematic **Fig. 3g**). Depletion of *Ddx6* significantly increased signal from *Mervl* and *Zscan4* reporters (**Fig. 3h, i**), suggesting that disrupting P-bodies facilitates the otherwise rare conversion of naïve ES cells to the 2C state. To validate that the disruption of P-bodies is indeed crucial for the phenotype resulting from *Ddx6* loss, we investigated whether inhibiting other essential factors for P-body assembly in ES cells could reproduce the phenotype observed with *Ddx6* suppression. Remarkably, knockout of *Eif4enif1* and *Lsm14a*^52^ led to the loss of P-bodies and activation of the *Mervl* and *Zscan4* reporters, akin to the effects observed in *Ddx6* depletion (**Fig. 3h, i**). These data support the hypothesis that P-bodies sequester fate-instructive 2C transcripts that are transiently suppressed and can enter translation upon P-body disassembly.

To explore this possibility further, we first assessed the integrity of P-body-associated transcripts in ES cells, including those encoding 2C regulators. Analysis of both SnapTotal-seq and SMART-seq data revealed over 70% overlap in P-body-associated transcripts (**Extended Data Fig. 5b**) between methods. Further comparison between P-body enrichment and mRNA half-lives in ES cells based on 4-sU labelling^90^ demonstrated minimal correlation between mRNA stability and P-body enrichment (r=-0.069) (**Extended Data Fig. 5c**). Additionally, we found no appreciable evidence of truncated mRNAs—indicative of degradation intermediates—or preferential deadenylation^91^ of P-body-associated RNAs compared to cytosolic fractions (r=-0.092) (**Extended Data Fig. 5d, e**), in line with previous reports^46, 49, 58^. These findings suggest that in ES cells, similar to our observations in HEK293T cells, P-body-associated transcripts are not preferentially degraded compared to cytosolic transcripts. We then examined the translational rate of RNAs within P-bodies of mouse ES cells. We compared our P-body-seq data with Ribo-seq data obtained in naïve and primed mouse ES cells^1,7^. Notably, we observed a strong, inverse correlation between P-body enrichment and translation efficiency suggesting that RNAs localized to P-bodies in naïve and primed ES cells are characterized by poor translation efficiency (**Fig. 4a**). These results indicate that RNAs in P-bodies in ES cells are intact and subject to translational suppression.

**Figure 4.**
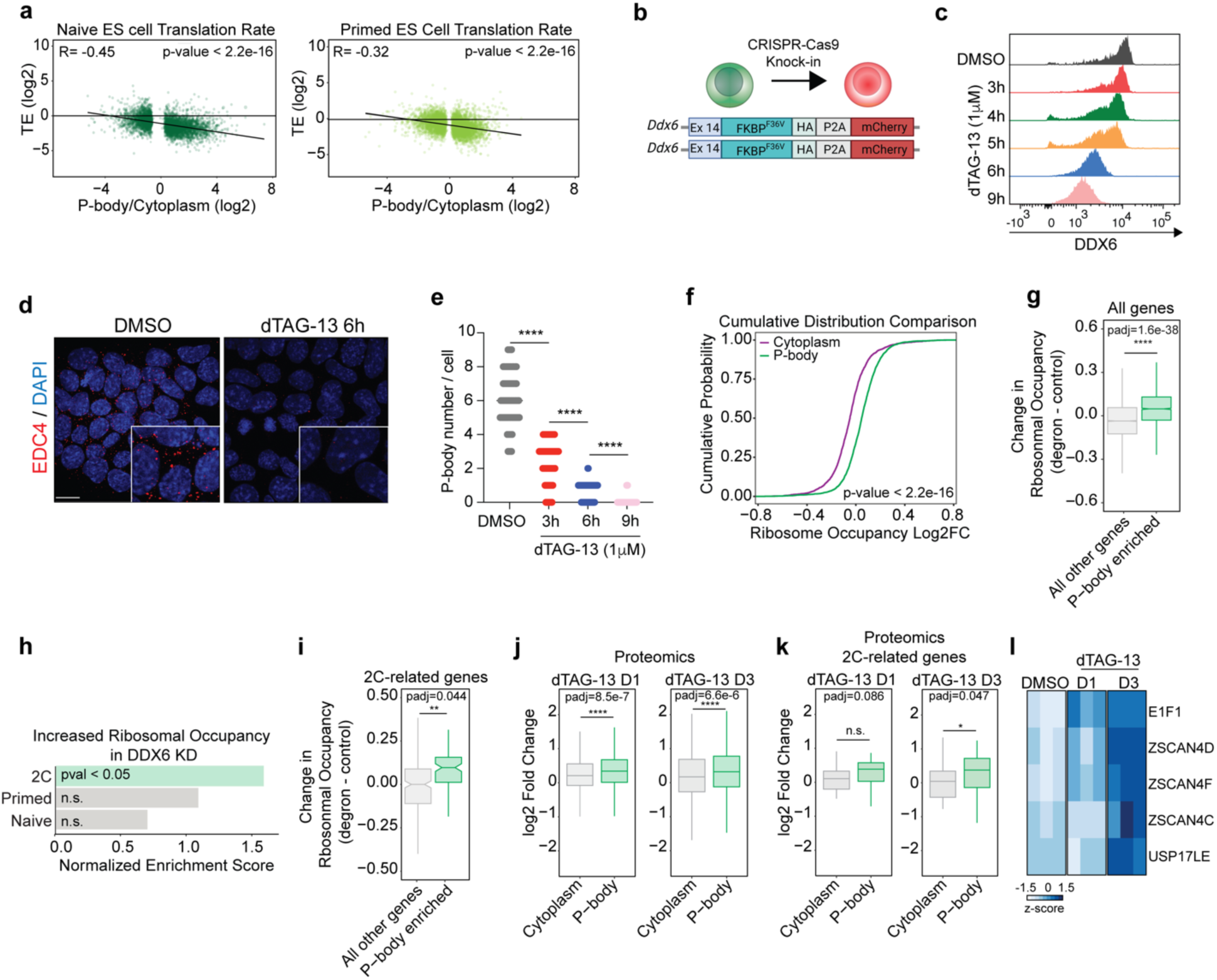
Transcripts related to the 2C state are sequestered in P-bodies, preventing their translation and activation of a 2C gene expression program. **(a)** Translation efficiency (log2 (Ribo-seq counts/RNA-seq counts)) negatively correlates with mRNA enrichment in P-bodies in naïve and primed mouse ES cells. Ribosome profiling data from^1^, Pearson correlation test. **(b)** A schematic showing CRISPR-Cas9-based homozygous insertion of FKBP12F36V-HA-P2A-mCherry sequence in place of the stop codon of the endogenous *Ddx6* allele. **(c)** Representative intracellular flow cytometry plots for DDX6 in *Ddx6*-FKBP12^F36V^ GFP-LSM14A mouse naïve ES cells, either untreated (DMSO) or treated with dTAG-13 at the indicated time points. **(d)** Representative IF imaging of EDC4 puncta (red) in *Ddx6*-FKBP12^F36V^ GFP-LSM14A mouse naïve ES cells, either untreated (DMSO) or treated with dTAG-13 for 6 hours. Nuclei were counterstained with DAPI (blue) (scale: 10μm). **(e)** P-body number in *Ddx6-*FKBP12^F36V^ GFP-LSM14A mouse naïve ES cells, either untreated (DMSO) or treated with dTAG-13 at the indicated time points. DMSO (n=70 cells), 3 hour-dTAG13 (n=70 cells), 6 hour-dTAG13 (n=70 cells), 9 hour-dTAG13 (n=70 cells), unpaired Student’s t-test, mean ± s.d., ****: p <0.0001. **(f)** Cumulative distribution function (CDF) plot showing ribosome occupancy (log2 FC) of P-body enriched and P-body-depleted mRNAs for untreated (DMSO) vs. dTAG-13 treated (6hrs) *Ddx6-*FKBP12^F36V^ GFP-LSM14A mouse naïve ES cells, ks-test. **(g)** Box plots showing the change in ribosome occupancy of all P-body enriched genes compared to all other genes. Unpaired t-test, mean ± s.d, padj (Holm’s method). **(h)** Normalized Enrichment Score (NES) of gene sets from (2C)^5^, (Naïve)^6^, and (Primed)^7^, p<0.05. **(i)** Box plots showing the change in ribosome occupancy of P-body enriched 2C-related genes^5^ compared to non-P-body enriched 2C genes. Unpaired t-test, mean ± s.d, padj (Holm’s method). **(j)** Box plots showing the change in protein levels of all P-body enriched genes compared to P-body depleted genes, after 1 day and 3 days of dTAG-13 treatment. Unpaired t-test, mean ± s.d, padj (Holm’s method) (* p < 0.05, ** p < 0.01, **** p < 0.0001) **(k)** Box plots showing the change in protein levels of P-body enriched 2C-related genes compared to P-body depleted genes, after 1 day and 3 days of dTAG-13 treatment. Unpaired t-test, mean ± s.d, padj (Holm’s method). **(l)** Heatmap showing protein levels of 2C-related genes after 1 day and 3 days of dTAG-13 treatment compared to control samples.

We next sought direct evidence that P-body-enriched mRNAs engage with translational machinery in stem cells following the acute dissolution of P-bodies. To achieve this, we generated a targeted DDX6 degron system in mouse naïve ES cells (*Ddx6*-FKBP12^F36V^) (**Fig. 4b**) and performed polysome profiling following acute loss of DDX6. Addition of dTAG-13, a potent and selective degrader of FKBP12^F36V^, facilitated DDX6 protein degradation and eliminated detectable P-bodies within 6 hours (**Fig. 4c-e and Extended Data Fig. 6a-c**). Consistent with our model, we observed a significant increase in the ribosome occupancy of P-body-associated mRNAs following DDX6 suppression compared to total cytosolic RNAs (p<2.2^−16^; **Fig. 4f, g**). Gene ontology analysis further supported that transcripts exhibiting enhanced ribosome occupancy were associated with a stem cell maintenance signature (**Extended Data Fig. 6d**). Given the observation that disrupting P-bodies in mouse naïve ES cells facilitated conversion to the 2C state, we next asked whether 2C-related transcripts demonstrated higher ribosome occupancy following DDX6 degradation. Indeed, 2C-associated mRNAs enriched in P-bodies exhibited a marked increase in ribosome occupancy after P-body dissolution (**Fig. 4h, i**). These data provide further evidence that mRNAs sequestered in P-bodies engage translational machinery upon acute loss of RNA sequestration in ES cells, which in turn, alters cell fate.

To determine whether increased translation of P-body-associated mRNAs led to corresponding protein-level changes, we performed large-scale, quantitative proteomics in *Ddx6* degron ES cells following dTAG-13 treatment for 1 and 3 days. This analysis revealed that P-body dissolution led to a significant upregulation of proteins encoded by P-body-associated mRNAs (**Fig. 4j**), including regulators of 2C-like cells (**Fig. 4k**). Elevated levels of these proteins facilitated the transition of ES cells to a 2C-like state, as shown by the increased abundance of totipotency factors such as ZSCAN4c/d/f, USP17LE, and EIF1^88^ (**Fig. 4l**). These findings demonstrate that disrupting RNA sequestration in P-bodies promotes the translation of proteins important for cell fate transitions.

In line with previous studies, a fraction of P-body-enriched RNAs underwent degradation following acute loss of DDX6 (**Extended Data Fig. 6e**). These downregulated mRNAs were linked to differentiation processes (**Extended Data Fig. 6f**), perhaps reflecting a separate mechanism for protecting transcripts needed for differentiation until appropriate signals/conditions are achieved during development^92^.

Collectively, our data underscore the role of P-bodies in restricting cell identity by buffering the activity of mRNAs related to earlier developmental stages through translational suppression. Furthermore, our findings thus far raise the intriguing possibility that a conserved mechanism drives mRNA sequestration into P-bodies in a context-dependent manner.

### miRNAs direct cell type-specific sequestration of mRNAs into P-bodies

Our data thus far suggest that mRNAs sequestered into P-bodies across various cell types encode specifiers of a previous developmental context (e.g., ZGA transcripts are enriched in P-bodies of naïve human ES cells). However, how this specificity is achieved was unclear. To address this point, we first considered the potential role of RNA modifications in directing transcripts into P-bodies. Among these modifications, the N^6^-methyladenosine (m^6^A) modification is the most abundant in eukaryotic mRNA^93^. To investigate whether m^6^A influenced RNA sequestration into P-bodies in pluripotent stem cells, we used a human ES cell model that permits doxycycline (dox)-inducible ablation of the m^6^A methyltransferase, METTL3 (TET-OFF *METTL3*)^94^ (**Extended Data Fig. 7a**). We confirmed that METTL3 was undetectable at both the protein and RNA levels after 8 days of dox treatment (**Extended Data Fig. 7b**), accompanied by a significant reduction in global m^6^A levels (**Extended Data Fig. 7c**). We further confirmed that genes related to pluripotency (i.e., *NANOG*) were unaffected by *METTL3* knockout (**Extended Data Fig. 7d**). Immunofluorescence for EDC4^+^ puncta revealed no change in P-body numbers upon METTL3 deletion (**Extended Data Fig. 7e**). We next profiled P-body content and found no significant differences in *METTL3* KO cells compared to control (85% overlap; **Extended Data Fig. 7f**). Moreover, intersecting the P-body transcriptome with a dataset of m^6^A-methylated mRNAs in human ES cells revealed that only a small fraction of P-body-enriched RNAs is m^6^A methylated and P-body enrichment for the majority of these transcripts did not change following *METTL3* KO (**Extended Data Fig. 7f**; compare white and gray circles). These results suggest that m^6^A RNA methylation is not primarily responsible for directing RNA sequestration into P-bodies in human pluripotent stem cells, which aligns with a recent report that shows negligible effects of m^6^A modification on mRNA recruitment to stress granules in mouse ES cells^95^.

Given the pivotal role of RBPs in post-transcriptional control of RNAs, we next explored their potential involvement in RNA sequestration within P-bodies. To address this point, we utilized published CLIP-seq data^96^ to map the P-body enrichment of transcripts targeted by 53 RBPs (**Fig. 5a**). Targets of proteins known to localize to RNA granules, including PUM2 and HNRNPU^38, 97^, were enriched in P-bodies; however, these proteins are widely expressed and unlikely to confer cell type-specificity. In line with this finding, de novo motif analysis of P-body-enriched transcripts predicted the recognition by multiple RBPs that were conserved across cell types, with no evident cell-type-specific patterns^98^ (**Extended Data Fig. 7g, h**). GO analysis of the identified RBPs revealed enrichment for pathways consistent with known P-body functions, including 3’ UTR processing, RNP-granule assembly, and mRNA stabilization (**Extended Data Fig. 7i**). Collectively, these data suggest that while RBPs play fundamental roles in P-body assembly and function, they likely do not drive the cell type-specific RNA sequestration patterns that we observed.

**Figure 5.**
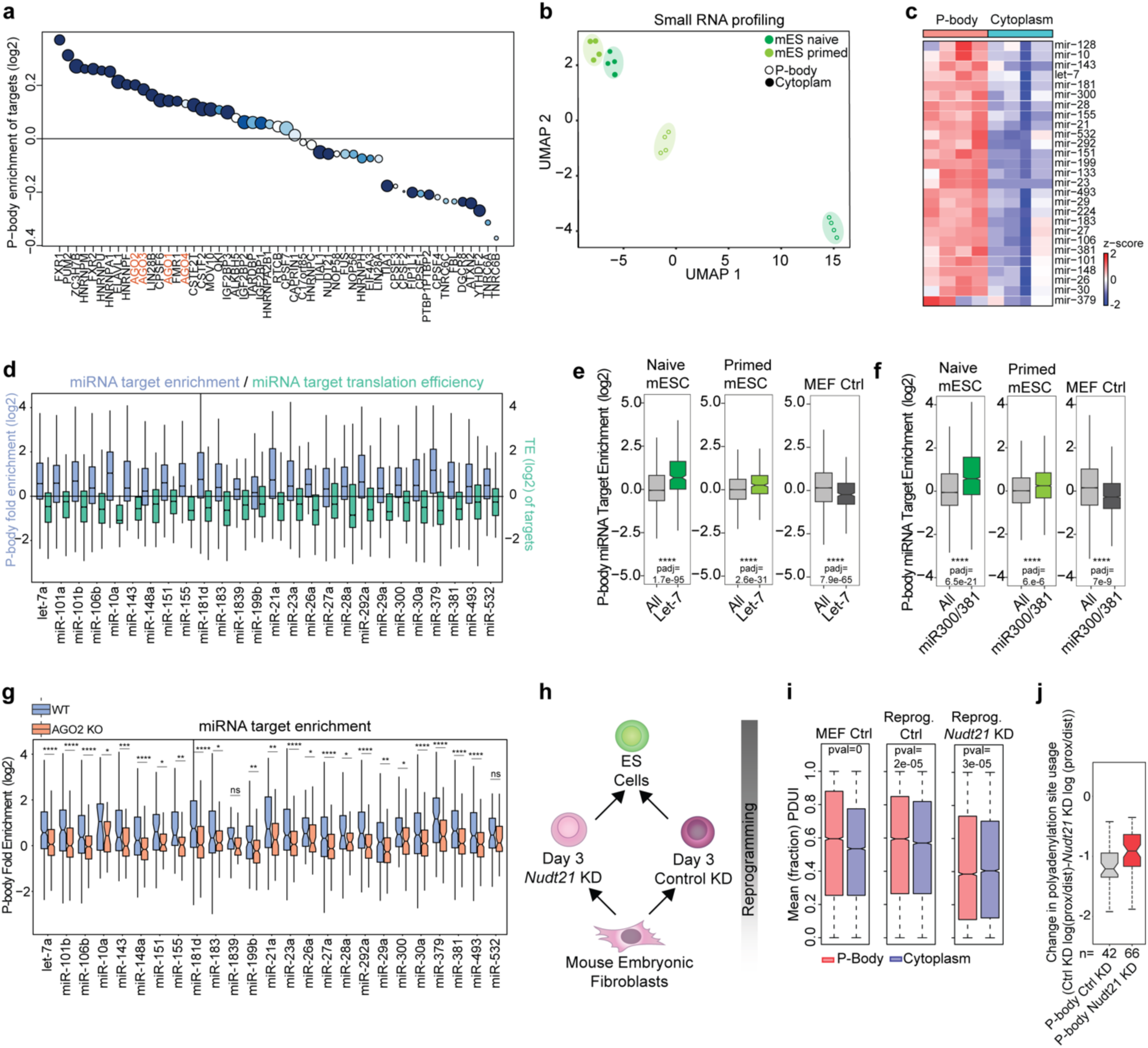
miRNAs regulate RNA sequestration in a context-dependent manner. **(a)** mRNA targets of 53 RNA-binding proteins were analyzed for their enrichment in P-bodies and cytoplasm, using CLIP-seq experiments available in the CLIPdb 1.0 database. Circle size indicates counts; color indicates p-value. **(b)** UMAP analysis of small RNA-seq data for the indicated samples based on differentially expressed genes between purified P-body and cytoplasmic fractions. **(c)** Heatmap showing expression levels of differentially enriched miRNAs between purified P-body and cytoplasmic fractions in mouse naïve ES cells. (n=2, p < 0.05). **(d)** Quantification of P-body enrichment for miRNA targets and their corresponding translation efficiency (TE). Ribosome profiling data from1. **(e, f)** Box plots showing P-body enrichment of Let-7 (e) and miR-300/381 targets (f) in naïve and primed mouse ES cells and mouse fibroblasts. Unpaired t-test ± s.d, showing Holm’s method adjusted p-values, ****p<0.0001. **(g)** Box plot comparing P-body enrichment of miRNA-targets between WT and AGO2 KO mouse naïve ES cells. Unpaired t-test, showing Holm’s method adjusted p-values, ns: p>0.05, *p<0.05, **p<0.01, ***p<0.001, ****p<0.0001. **(h)** Schematic of iPS cell reprogramming, including perturbation of the alternative polyadenylation regulator Nudt21. **(i)** PolyA site usage index (PDUI) of p-body and cytoplasm enriched transcripts. PDUI<0.5 indicates proximal polyA preference, PDUI>0.5 indicates distal polyA preference Unpaired t-test, showing Holm’s method adjusted p-values. **(j)** Change in polyadenylation site usage upon Nudt21 KD.

Further inspection of the CLIP-seq data revealed enrichment of targets associated with members of the Argonaute protein family (AGO1, AGO2, AGO3, and AGO4) within P-bodies (**Fig. 5a**). AGO proteins serve as essential components of the RNA-induced silencing complex (RISC) and cooperate with miRNAs to suppress the translation of mRNAs^99, 100^. Previous studies have shown localization of AGO proteins in P-bodies and suggested a connection between certain miRNAs and mRNA recruitment to RNP granules in non-mammalian cells and transformed cell lines^42, 101–106^. Moreover, miRNAs are known to be expressed in a cell type-specific manner and play a prominent role in cell fate transitions^107–110^. These intriguing observations led us to hypothesize that miRNAs may play a pivotal role in directing RNA sequestration of developmental transcripts into

#### P-bodies in a cell type-specific manner

To test this hypothesis, we performed comprehensive profiling of small RNA populations in both P-body and cytoplasmic fractions across mouse and human pluripotent cells and neural progenitors using small non-coding RNA-seq^111^ (**Extended Data Fig. 8a**). Uniform Manifold Approximation and Projection (UMAP) plot analysis revealed clustering of naïve and primed mouse ES cell samples according to their cellular origin (**Fig. 5b**). Remarkably, our analysis revealed that subsets of miRNAs were selectively enriched in P-bodies of pluripotent cells as well as progenitor cells (**Fig. 5c and Extended Data Fig. 8b, c**). For example, among others, we identified miR-300 and miR-Let-7, which are known regulators of stem cell potency^112, 113^. In line with our hypothesis, targets of miRNAs enriched in P-bodies were also sequestered within these RNA condensates (**Fig. 5d and Extended Data Fig. 8d, e**) and, importantly, are translationally repressed (**Fig. 5d**). Supporting these data, analysis using MicroRNA ENrichment TURned NETwork (MIENTURNET)^114^ revealed that P-body-targeted transcripts were enriched for specific miRNA binding sites that were absent in cytoplasmic transcripts (**Extended Data Fig. 8f-h**).To explore this relationship further, we plotted the P-body enrichment for targets of miR-Let-7 and miR-300/381 (miR-300 and miR-381 share a highly similar target sequence) across cell types. In both naïve and primed ES cells, we observed an increased enrichment of their targets in P-bodies (**Fig. 5e, f**). By contrast, in mouse embryonic fibroblasts (MEFs), we found no enrichment for target transcripts in P-bodies (**Fig. 5e, f**). Together, our data suggest that miRNA-targeted genes are sequestered into P-bodies in a cell-type specific manner.

### Disrupting miRNA function prevents RNA sequestration in P-bodies

Our data thus far suggest that miRNAs target mRNAs for storage in P-bodies across cell types. To provide functional evidence for this relationship, we sought to uncouple miRNA interactions with target mRNAs. Previous studies demonstrated the importance of AGO2 in the sequestration of Let7-targets into P-bodies in transformed cells^101^. To investigate the involvement of AGO2 in global RNA sequestration in pluripotent stem cells, we generated AGO2 knockout (KO) mouse ES cells using CRISPR-Cas9^115^ (**Extended Data Fig. 9a**). AGO2 KO cells exhibited a similar number of P-bodies compared to WT cells, as indicated by the presence of EDC4^+^ puncta (**Extended Data Fig. 9b**). This finding confirms that AGO2 knockout does not prevent formation or maintenance of P-bodies. To assess the subcellular localization of miRNA-target transcripts following AGO2 deletion, we sorted P-bodies from AGO2 KO ES cells and compared their transcriptome to that of their WT counterparts. We found that AGO2 KO significantly reduced miRNA-targeted mRNAs in P-bodies (**Fig. 5g**).

Next, we investigated whether the modulation of polyA site usage (i.e., Alternative Polyadenylation (APA)) influenced RNA sequestration into P-bodies. APA generates RNA isoforms that commonly differ by the length of their 3’ UTR^116–119^. Because miRNA target sequences are frequently found in the 3’ UTR, we reasoned that forcing proximal polyadenylation (i.e., shorter 3’ UTRs) would relieve miRNA-based targeting of transcripts into P-bodies. To test this possibility, we suppressed expression of *Nudt21*, a component of the CFIm complex that facilitates distal polyadenylation^18, 117^, in MEFs during reprogramming to pluripotency (**Fig. 5h**). We chose this context because a matched dataset of APA changes is available^18^, permitting direct comparison between polyA site usage and P-body enrichment. Of note, *Nudt21* knockdown leads to increased reprogramming efficiency and expression of pluripotency-related genes^18^, which we confirmed in our system at both the protein and RNA levels (**Extended Data Fig. 9c, d**). Similar to AGO2 knockout, the presence and number of P-bodies were not significantly affected by *Nudt21* suppression (**Extended Data Fig. 9e**). We mapped the P-body transcriptome of uninduced MEFs as well as reprogramming intermediates with and without *Nudt21* silencing (3 days of reprogramming induction). We observed that in MEFs and day 3 control reprogramming cells, 3’ UTRs of P-body-targeted RNAs were longer compared those of their cytoplasmic counterparts, consistent with a role for regulatory sequences within 3’ UTRs in mediating P-body enrichment (**Fig. 5i**). Following *Nudt21* suppression, we observed a global reduction in 3’ UTR length and the 3’ UTRs of transcripts enriched in P-bodies had similar length relative to cytoplasmic transcripts (**Fig. 5i**). We next used published data to compare changes in polyadenylation site usage to changes in transcript localization between P-bodies and cytoplasm following *Nudt21* knockdown^18^. Notably, transcripts that were enriched in the P-bodies of control cells, but were not identified in the P-bodies of *Nudt21* knockdown cells, had substantially shorter 3’ UTRs following *Nudt21* suppression (**Fig. 5j**). These results suggest that the loss of miRNA binding sites in the 3’ UTR regions of mRNAs prevents their localization to P-bodies. Collectively, these findings support the hypothesis that RNA sequestration in P-bodies is mediated by the RNA silencing machinery.

### Modulating miRNA function relocalizes their target transcripts and influences cell identity

Given the critical role of miRNAs in directing RNA sequestration in P-bodies, we next examined whether manipulating specific miRNAs could impact RNA localization and cell identity. To this end, we depleted miR-300 in mouse ES cells using an antisense inhibitor (**Fig. 6a**). Following 48 hours of anti-miR-300 treatment, we assessed the P-body localization of miR-300 target transcripts. In line with our AGO2 KO studies, we observed a significant reduction of miR-300 targets in P-bodies upon miRNA inhibition (**Fig. 6b**).

**Figure 6.**
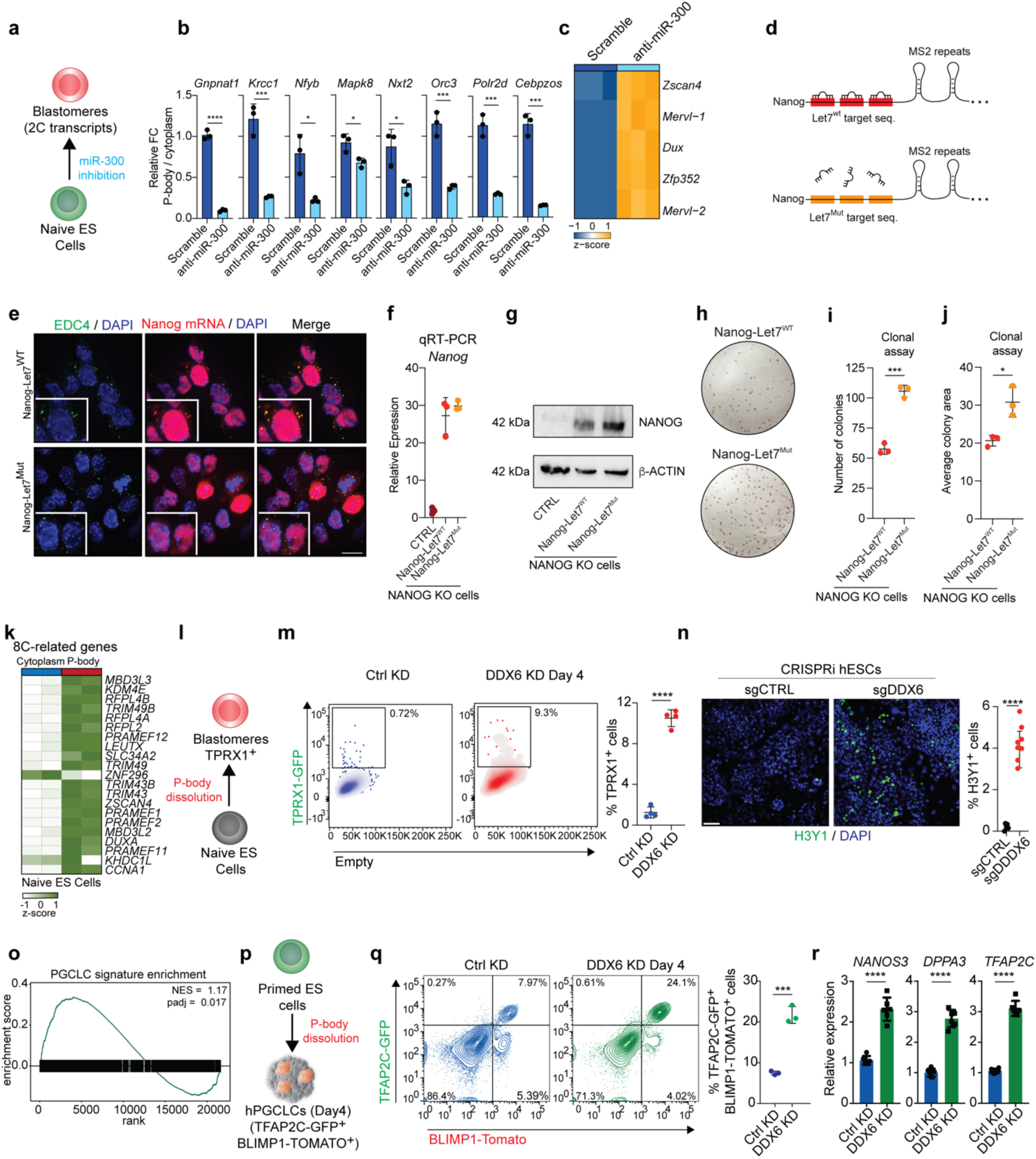
Modulating of miRNA function and P-body assembly alters cell fate. **(a)** A schematic of the strategy for miR-300 inhibition in mouse naïve ES cells. **(b)** qRT-PCR analysis of the expression for miR-300 targets in P-bodies of mouse naïve ES cells after miR-300 inhibition and for control. n=3, unpaired Student’s t-test, mean ± s.d., *p<0.05, **p<0.01, ****p<0.0001. **(c)** Heatmap showing expression levels of differentially enriched 2C-related genes in mouse naïve ES cells after miR-300 inhibition and for control. **(d)** A schematic of the strategy for the generation of Let7^wt^ and Let7^mut^ reporter cell lines. **(e)** Representative IF imaging of EDC4 puncta (green), and Nanog-MS2 (red) in Nanog Let7^wt^ (upper panel) and Nanog Let7^mut^ cells (lower panel). Nuclei were counterstained with DAPI (blue) (scale: 10μm). **(f)** qRT-PCR analysis of Nanog expression in Nanog Let7^wt^ and Nanog Let7^mut^ cells compared to CTRL cells (Nanog KO). **(g)** Representative western blot showing NANOG protein levels in Nanog Let7^wt^ and Nanog Let7^mut^ cells compared to CTRL cells (*Nanog* KO). **(h-j)** Representative pictures (h) and quantification of Alkaline Phosphatase staining of cell colony number (i) and size (j) from Nanog Let7^wt^ and Nanog Let7^mut^ cells cultured in FBS+LIF conditions. **(k)** Heatmap showing expression levels of differentially enriched 8C-related mRNAs between purified P-body and cytoplasmic fractions in human naïve ES cells. (n=2). **(l)** A schematic of the strategy for P-body dissolution in naïve human ES cells carrying blastomere-specific reporter (TPRX1-GFP). **(m)** Quantification of TPRX1-GFP^+^ cells upon *DDX6* KD. Unpaired Student’s t-test, control (n=3), *DD*X6 KD (n=3), mean ± s.d., ****: p<0.0001. **(n)** Representative IF imaging of H3Y1 (green) positive cells in naïve human ES cells upon *DDX6* KO compared to control cells. Nuclei were counterstained with DAPI (blue) (scale: 50μm) (Left panel). Quantification of H3Y1^+^ cells upon *DDX6* suppression. Unpaired Student’s t-test, sgControl (n=5 fields), sgDDX6 (n=8 fields), mean ± s.d., ****: p<0.0001 (Right panel). **(o)** GSEA analysis for hPGCLC-related gene expression signature in purified P-bodies from human primed ES cells. **(p)** A schematic of hiPSC to PGCLC differentiation. HiMeLCs: human incipient mesoderm-like cells. **(q)** Flow cytometric analysis of TFAP2C-GFP and BLIMP1-TOMATO expression after four days of PGCLC differentiation in 3D aggregates (left). Quantification of TFAP2C-GFP^+^/BLIMP1-TOMATO^+^ cells by flow cytometry for three experiments (right). Error bars indicate mean ± s.d. (*n=3*), statistical significance was determined using a two-tailed unpaired Student’s t-test (***P<0.001). **(r)** Quantitative RT-PCR analysis for the indicated genes after four days of PGCLC differentiation in 3D aggregates. Error bars indicate mean ± s.d. (*n>3*), statistical significance was determined using a two-tailed unpaired Student’s t-test (****P<0.0001).

To test whether releasing miR-300 targets from P-bodies was sufficient to induce a functional cell fate change (**Fig. 6a**), we examined the expression of 2C-related genes. Indeed, we observed robust upregulation of 2C-related transcripts, including *Zscan-4* and *Mervl*, upon miR-300 suppression, suggesting that miR-300 restricts reversion of pluripotent cells to an earlier developmental fate (**Fig. 6c**). Notably, these 2C transcripts are not direct targets of miR-300, suggesting that their upregulation results from cell fate change, rather than regulation from the miRNA itself. These findings support the hypothesis that RNA sequestration in P-bodies can be precisely manipulated to change cell identity.

We next tested whether it was possible to direct mRNA transcripts into P-bodies through the addition of miRNA-target sequences. To address this, we adapted reporter constructs based on the pluripotency master regulator, *Nanog*. The reporter constructs included a 3’ UTR with either six wild-type or six mutant Let-7 sequences (Let-7^WT^ and Let-7^Mut^), which have been previously used to test miRNA function^120, 121^, and 24 MS2 stem loops (**Fig. 6d**). These stem loops recruit a tagged version of the MS2 Coat Protein (JF546 HaloTag-MCP) to permit direct, subcellular visualization of *Nanog* mRNA. Following transfection into *Nanog* knockout ES cells, *Nanog-*Let-7^WT^-MS2 transcripts colocalized with the P-body marker EDC4 (**Fig. 6e**), suggesting that Let-7 target sequences were sufficient to direct *Nanog* mRNA into P-bodies. By contrast, *Nanog-*Let-7^Mut^-MS2 transcripts showed no appreciable accumulation in P-bodies (**Fig. 6e**). Importantly, both constructs produced similar steady state RNA levels (**Fig. 6f**), excluding the possibility that the observed differences between Let-7^WT^ and Let-7^Mut^ arise from differences in RNA abundance. NANOG protein levels, however, were lower in cells expressing *Nanog-*Let-7^WT^-MS2, supporting a non-degradative, post-transcriptional regulatory mechanism (**Fig. 6g**).

Finally, we investigated whether miRNA-dependent recruitment of RNAs to P-bodies could be leveraged to alter cell identity. Under naïve conditions, *Nanog* knockout cells are capable of self-renewal; however, when switched to standard ES culture media (i.e., FBS with LIF), NANOG plays a crucial role in sustaining self-renewal. We therefore transitioned *Nanog* knockout cells expressing either *Nanog-*Let-7^WT^-MS2 or *Nanog-*Let-7^Mut^-MS2 from naïve to standard culture conditions. Cells expressing Nanog-MS2-Let-7^WT^ had substantially lower self-renewal capacity compared to *Nanog-*Let-7^Mut^-MS2 expressing cells, as evidenced by increased colony number and size in clonal assays (**Fig. 6h-j**). These data highlight the crucial role of miRNA-mediated RNA sequestration in regulating cell potency. Altogether, these findings underscore the critical role of miRNA-mediated RNA sequestration in regulating cell potency and demonstrate that actively directing mRNAs to P-bodies can be exploited to modulate cell identity.

### Manipulating P-body-mediated RNA sequestration to direct cell fate

Our data identify RNA sequestration in P-bodies as a fundamental regulator of cell fate specification. We therefore sought to leverage this mechanism to facilitate inefficient cell fate conversions from human pluripotent stem cells to rare, clinically relevant cell types. We first focused on the generation of human totipotent-like cells, which remain difficult to obtain *in vitro* at high efficiency. Thus far, totipotent-like cells have been identified *in vitro* at a 0.1% frequency in naïve pluripotent cells through single-cell RNA-seq^3, 122^. Our sequencing-based analysis of naïve human pluripotent stem cells revealed that P-bodies sequester transcripts associated with the totipotent 8-cell stage embryo (**Fig. 2i and 6k**), likely preventing their translation and restricting naïve ES cell plasticity. To test whether P-body dissolution could promote the conversion of human naïve ES cells to a totipotent-like state, we generated a human naïve ES cell line carrying a totipotency-specific reporter, TPRX1-GFP^83^ (schematic in **Fig. 6l**). Remarkably, DDX6 suppression increased the proportion of TPRX1-GFP^+^ cells by 100-fold (∼0.1% to ∼10%) under naïve conditions over previously reported efficiencies^3^ (**Fig. 6m**). To further validate these findings, we employed an orthogonal CRISPRi-based approach to deplete *DDX6*^48^, which led to the activation of *H3Y1*, a DUX4 target gene expressed in the 8-cell embryo^123^(**Fig. 6n**). Together, these findings provide functional evidence that P-bodies sequester totipotency-associated transcripts to suppress their expression. Disrupting this key regulatory module is sufficient to transition naïve ES cells to a totipotent-like state. Moreover, our data indicate that P-bodies might be exploited to induce and expand rare human cell types *in vitro*.

We next explored whether modulating P-bodies could enhance the generation of specialized cell types from human pluripotent stem cells. We focused on the conversion from human ES cells to primordial germ cell-like cells (PGCLCs) because they are difficult to derive *in vitro,* share epigenetic and transcriptional features with totipotent stem cells^124, 125^, and are highly relevant for reproductive medicine. Moreover, our data indicated that P-bodies in primed human ES cells sequester transcripts associated with a primordial germ cell signature (**Fig. 6o**), including *FXR1*^126^, *ING2*^127^, *EIF4G3*^128^, and *MEIOC*^129^(**Extended Data Fig. 9f**). Based on these observations, we hypothesized that modulating P-bodies could enhance the programming of primed human ES cells toward the germ cell lineage. To test this, we introduced shRNAs targeting *DDX6* into primed human pluripotent cells carrying the PGC-specific reporters, TFAP2C-GFP and BLIMP1-TOMATO^130^ (**Fig. 6p**). Immediately following DDX6 knockdown, we induced germ cell fate using a multi-step differentiation protocol^130^. In our hands, we achieved a maximum conversion efficiency of less than 8% in standard conditions; however, *DDX6* suppression consistently increased the formation of TFAP2C/BLIMP1 double-positive PGCLCs by more than threefold to greater than 24% (**Fig. 6q**). Furthermore, DDX6-depleted aggregates expressed significantly higher levels of key PGC markers, including NANOS3, DPPA3 and TFAP2C (**Fig. 6r**). These findings demonstrate that P-body dissolution facilitates the efficient programming of primed human ES cells toward the germ cell lineage. Collectively, our findings establish a framework for harnessing condensate biology to expand clinically relevant stem cell populations, highlighting P-body modulation as a promising strategy for cell fate engineering.

## DISCUSSION

Development and homeostasis require rapid and precise regulation of fate-instructive genes to direct cell specification. While the role of transcriptional regulation in controlling cell fate decisions is widely appreciated, our understanding of posttranscriptional mechanisms is still emerging. Here, we identified RNA sequestration through P-bodies as a fundamental regulatory mechanism in development and differentiation (**Extended Data Fig. 9g**). By leveraging fluorescence-activated particle sorting, we directly compared P-body contents between cell types at distinct developmental stages. Our analyses revealed cell type-specific RNA sequestration patterns that reflected transcriptional profiles characteristic of preceding developmental stages. For example, important totipotency-associated transcripts were directed into P-bodies in mouse and human naïve ES cells, while mRNAs for neural progenitor-related genes were enriched in the P-bodies of neurons. These data suggest that transcripts important for stem and progenitor cells are specifically directed to P-bodies to inhibit their translation upon differentiation. In support of this model, disrupting P-bodies in naïve mouse ES cells led to the upregulation of established 2C reporters and drove a gene expression signature characteristic of the 2C state. Notably, we observed increased ribosome occupancy and protein levels for P-body-enriched transcripts, including 2C genes, following acute disruption of P-bodies in mouse naïve ES cells, suggesting that translation of these transcripts is directly inhibited by their sequestration. Together, these data provide functional evidence that P-body-based regulation is sufficient to alter cell fate.

Our findings underscore the potential of manipulating P-body-mediated regulation to facilitate the conversion of pluripotent stem cells into clinically relevant cell types, including totipotent-like cells. While human totipotent stem cells have previously been identified at a low frequency (∼0.1%) in naïve pluripotent cell populations using single-cell RNA sequencing^3^ or generated using chemical reprogramming from primed cells—via inhibition of four distinct pathways—at low efficiency (∼1%)^83^, our data demonstrate that disrupting P-body-mediated regulation dramatically enhances the frequency of TPRX1-GFP^+^ totipotent-like cells to nearly 10% under naïve culture conditions. Totipotent cells hold promise not only for producing high-quality blastoids but may also enable the generation of differentiated cells and tissues with unprecedented efficiency^83, 122, 131, 132^. Furthermore, mouse totipotent cells express transcription factors absent in humans^133, 134^, suggesting that the robust expansion of human totipotent cells we achieved could provide a unique platform to investigate species-specific regulatory mechanisms governing early mammalian embryogenesis. In addition to totipotent-like cells, we show that RNA condensates can be leveraged to promote the specification of human pluripotent stem cells into PGCLCs. The development of systems to generate large quantities of PGCLCs is invaluable to further our understanding of germ cell specification and for developing therapeutics for infertility^125, 130, 135, 136^. Based on these proof-of-principle experiments, we propose that strategies to selectively mobilize specific mRNAs from P-bodies, including the use of antisense oligonucleotides, could offer a novel approach for precise cell fate engineering. Moreover, our P-body-seq has identified fate-instructive factors within P-bodies in adult progenitor cells of the three germ layers, raising the intriguing possibility that manipulating P-bodies in these cells could similarly redirect cell fate or expand cell populations that are currently challenging to propagate. Collectively, these insights establish a foundation for leveraging condensate biology to engineer cell fate, with broad implications for regenerative medicine, developmental biology, and therapeutic innovation.

A fundamental question emerging from our findings is why differentiating cells employ RNA sequestration rather than degradation to suppress the expression of transcripts related to stem and progenitor cell self-renewal. We propose that RNA sequestration represents a reversible regulatory mechanism that provides cells with developmental plasticity during fate transitions. The temporary storage of stem cell transcripts provides cells with an opportunity to revert to a self-renewal program if environmental conditions prove unfavorable for differentiation. This model is analogous to the “gatekeeping” function of NANOG^137^, which is transiently downregulated in a subset of ES cells under self-renewing conditions. Lowering levels of NANOG is thought to increase the propensity for cells to differentiate, but NANOG expression can reinitiate in the absence of differentiation cues to safeguard pluripotency. We speculate that a similar mechanism, regulated by P-body localization, permits stem and progenitor cells to sample the environment and direct cell fate accordingly. Supporting this model, we find that sequestered transcripts ultimately decline in terminally differentiated cells, suggesting that P-body localization serves as a key regulatory mechanism during processes that require rapid and coordinated changes in gene expression programs, such as development and stress responses. The transient sequestration of developmental stage-specific mRNAs in P-bodies may provide cells with developmental flexibility during early differentiation, while this mechanism becomes dispensable once cells achieve terminal differentiation and establish stable gene expression programs for tissue homeostasis. Interestingly, this pattern parallels the role of several miRNAs, which are essential for developmental transitions but less critical for adult tissue maintenance^138, 139^, further supporting our model of miRNA-based RNA sequestration.

To provide further evidence that RNA sequestration is a general regulatory mechanism for regulating cell fate, we asked whether P-body enrichment of specific transcripts was conserved through evolution. Indeed, comparative analyses between mouse, human, and chicken ES cells revealed substantial overlap in P-body constituents between species. Moreover, both mouse and human naïve ES cells transitioned to a totipotent-like state following disruption of P-bodies, suggesting functional conservation. The observation that P-body based regulation is conserved across multiple species supports RNA sequestration as a fundamental mechanism for controlling gene expression and cell fate. Whether P-body-based regulation controls cell fate in more distant organisms remains an open question.

The observation that P-bodies store distinct transcripts in diverse developmental contexts raises the question of how particular mRNAs are selected for sequestration. Within P-bodies, we found enrichment for targets of RNA-binding proteins, including known P-body components such as PUM2 and multiple Argonaute proteins. Since RNA-binding proteins like PUM2 are widely expressed, they are unlikely to direct cell type-specific sequestration of transcripts. However, while Argonaute proteins are also widely expressed, the miRNAs that guide them to target mRNAs are often cell type-specific. Correspondingly, we show that a subset of miRNAs is enriched in P-bodies in a cell type-specific manner. Disrupting miRNA-target interaction through multiple, orthogonal approaches prevented enrichment of corresponding miRNA targets in P-bodies. These findings suggest that the interplay between cell type-specific miRNA expression levels and the available repertoire of mRNA targets within a given cellular context is a key determinant of P-body composition. It remains unclear, however, whether this context-dependence arises purely from the action of distinct miRNAs in different cell types, or if common miRNAs mediate the sequestration of specific transcripts based on their abundance and accessibility within different cell contexts. Additional factors, including miRNA target abundance, 3’ UTR length, and the presence or absence of additional regulators likely contribute to cell-type specific P-body enrichment. Regardless, our data demonstrate that miRNA-mRNA interactions are a critical mechanism regulating P-body composition, revealing a previously unappreciated mechanism whereby miRNAs direct specific transcripts to biomolecular condensates for translational suppression and storage, rather than degradation. Future studies clarifying how RNA sequence, abundance, structure, and interaction partners influence the properties of condensates will be crucial for understanding their diverse context-specific roles in regulating cell fate decisions.

Overall, our data are consistent with specific sequestration of cell fate-instructive mRNAs during development. These findings extend our understanding of P-body-based regulation and add an additional layer of nuance to the mechanisms controlling cell fate decisions. We recognize that miRNAs likely drive sequestration of some transcripts, but not all. Additional mechanisms that facilitate localization of mRNAs into P-bodies remain an interesting avenue for future research. Beyond advancing our understanding of condensate function in cell fate control, our findings have potential therapeutic implications, as modulating P-body dynamics could provide a strategy to expand clinically relevant cell populations.

## ACKNOWLEDGMENTS

We thank all members of the Brumbaugh and Di Stefano groups for stimulating scientific discussions. We are grateful to Miguel Esteban for the TPRX1-GFP construct, Tim J. Stasevich for helpful discussions and MS2 constructs, Kyle Loh for the help with the endoderm differentiation, and Marc Fabian for the Let-7 constructs. B.D.S. is a Cancer Prevention and Research Institute of Texas (CPRIT) Scholar in Cancer Research. B.D.S. is supported by the CPRIT Recruitment of First-Time, Tenure-Track Faculty Member Award RR200079, the American Society of Hematology Scholar Award, the Milky Way Research Foundation Investigator Award, the Nancy Chang, Ph.D. Award for Research Excellence, the B+ Foundation, the Worldwide Cancer Research Foundation, and the NIH MIRA award 1R35GM147126. J.B. is grateful for support from the National Institutes of Health (R35GM142884) and Boettcher Foundation’s Webb-Waring Biomedical Research Awards program. M.N. was funded by the National Institutes of Health (T32GM142607). S.K. was funded by the NIH award F32 CA288043-01. E.J.P. was supported by the NIH award 1F30HD114315. Imaging for this project was supported by the Integrated Microscopy Core at Baylor College of Medicine and the Center for Advanced Microscopy and Image Informatics (CAMII) with funding from NIH (DK56338, CA125123, ES030285, S10OD030414), and CPRIT (RP150578, RP170719). This project was supported by the Cytometry and Cell Sorting Core at Baylor College of Medicine with funding from the CPRIT Core Facility Support Award (CPRIT-RP180672), the NIH (CA125123 and RR024574). NIH funding (P01 HL146366) to N.J.K.

## AUTHOR CONTRIBUTIONS

P.P., M.N., J.B., and B.D.S. conceived the study and wrote the manuscript; J.S. and Q.C. conducted and analyzed the PANDORA-seq and small-RNA-seq data; S.K. carried out the FAPS analysis; Y.C. created the 8C human ES reporter line; A.L.R., E.S. and D.L.S. conducted the Proteomics; E.C. contributed to the bioinformatic analysis; N.J.K. funding acquisition; E.J.P. conducted the reprogramming experiments; X.C. and Q.Y. derived and expanded the chicken ES cells; F.L.F. performed TET-OFF *METTL3* cell culture experiments.

## METHODS

### Mouse embryonic stem cell culture

Mouse ES cells used in this study were C57BL/6 x 129S4Sv/Jae F1-derived V6.5 ES cells. These cells were maintained at 37°C in a culture medium suitable for naïve mouse ES cell, which consisted of a 1:1 ratio of DMEM/F12 medium (Sigma Aldrich) and Neurobasal medium (Life Technologies). The culture medium was supplemented with 1X MEM non-essential Amino Acid Solution (Sigma-Aldrich), 1 mM Sodium Pyruvate (Sigma-Aldrich), 2 mM L-Glutamine (Sigma-Aldrich), 100 U/mL Penicillin (Sigma-Aldrich), 100 mg/mL Streptomycin (Sigma Aldrich), 50 μM β-mercaptoethanol (Sigma-Aldrich), and N2 and B27 supplements (referred to as N2B27 medium), two small-molecule inhibitors PD0325901 (1 μM, Axon Medchem) and CHIR99021 (3 μM, Axon Medchem), and Leukemia Inhibitory Factor (mLIF) (10 ng/mL, R&D Systems).

For the induction of mouse primed ES cells, a seeding density of 1.0×10^5^ mouse naïve ES cells was used. These cells were plated onto the wells of a 12-well plate, which had been pre-coated with Matrigel (Corning). The culture medium for this induction consisted of N2B27 medium supplemented with 20 ng/mL of Activin A (Peprotech), 12 ng/mL of bFGF (Peprotech), and 1% KnockOut Serum Replacement (KSR, Life Technologies).

### Derivation of mouse embryonic fibroblasts

Mouse embryonic fibroblasts (MEFs) were generated following a protocol similar to that described by^140^. Briefly, embryos were collected from timed mating at embryonic day 13.5 (E13.5). Subsequently, the head, limbs, spinal cord, gonads, and internal organs were carefully excised, leaving behind the remaining tissue. This tissue was then finely minced and treated with 200 μL of Trypsin-EDTA (Sigma-Aldrich). After a 5-minute incubation at 37°C, the enzymatic reaction was halted by adding 10 mL of MEF growth media composed of DMEM (Sigma-Aldrich), supplemented with 10% fetal bovine serum (Corning), L-Glutamine (2 mM, Sigma-Aldrich), and penicillin-streptomycin (Sigma-Aldrich). The cells were cultured at 37°C under low oxygen conditions (5% O2).

For reprogramming experiments, MEFs were isolated from mice harboring an inducible, polycistronic OKSM cassette located in the 3’ untranslated region (UTR) of the *Col1a1* gene. Additionally, these MEFs carried the M2-rtTA transactivator at the Rosa26 locus, along with an EGFP reporter construct regulated by *Pou5f1* regulatory elements^141^. All procedures involving mice were conducted in accordance with the guidelines stipulated by our Institutional Animal Care and Use Committee (IACUC) and were in compliance with the approved protocol no. AN-8464.

### Induction of pluripotency

To initiate the expression of pluripotency factors, reprogrammable MEFs were cultivated in induction media containing KO-DMEM (Life Technologies), with 15% fetal bovine serum (Corning), 1X non-essential amino acids (Life Technologies), L-Glutamine (2 mM, Sigma-Aldrich), 1000 U/ml LIF (10 ng/mL, R&D Systems), 55 mM beta-mercaptoethanol (Sigma-Aldrich), and supplemented with 50 μg/ml ascorbic acid (Sigma Aldrich) and 2 μg/ml doxycycline hyclate (Sigma-Aldrich).

For P-body isolation, reprogrammable MEFs were subjected to induction for a duration of 3 days, at which time, samples were harvested. Alternatively, to achieve complete reprogramming, these MEFs were maintained under induction conditions for a period of 8 days.

### Human primed pluripotent stem cell culture and naïve conversion

Conventional human primed ES cells (UCLA-4 and WIBR3 “TETOFF METTL3”, both female cell lines) were cultivated on Matrigel-coated dishes in mTeSR1 medium (Stem Cell Technologies) at a temperature of 37°C. These cells were propagated by using a 0.02% EDTA solution (Sigma-Aldrich) for passaging purposes. Cell passaging occurred every 4-5 days to ensure proper maintenance.

In the context of the doxycycline experiment, the TETOFF METTL3 cell line was cultured in mTeSR1 Medium supplemented with 2 μg/ml doxycycline (Sigma-Aldrich) for a duration of 1 week.

To achieve reversion to a naïve state, female UCLA-4 human ES cells that had been passaged six days prior underwent a procedure involving washing with 1X PBS (Life Technologies) and then treating with TrypLE Express enzyme (1X, Life Technologies) for a period of 3 minutes. This enzymatic treatment enabled cell dissociation into a single-cell suspension, which was subsequently plated at a density of 30,000 cells per 9.5cm² on irradiated CF-1 MEFs (a mixture of pooled male and female cells). This process occurred in human ES cell medium supplemented with 10 μM Y-27632 (Axon Medchem). Following a two-day incubation period, the medium was changed to 5i/LAF and then replaced daily. The 5i/LAF medium composition consisted of a 50:50 mixture of DMEM/F-12 (Sigma-Aldrich) and Neurobasal medium (Life Technologies), supplemented with 1X N2 supplement (Life Technologies), 1X B27 supplement (Life Technologies), 10 ng/mL bFGF (Peprotech), 1% non-essential amino acids (Sigma-Aldrich), L-Glutamine (2 mM, Sigma-Aldrich), penicillin-streptomycin (Sigma-Aldrich), 0.1 mM beta-mercaptoethanol (Sigma-Aldrich), 50 μg/mL BSA (Sigma-Aldrich), 0.5 μM IM-12 (Axon Medchem), 0.5 μM SB590885 (Axon Medchem), 1 μM WH-4-023 (Axon Medchem), 10 μM Y-27632 (Axon Medchem), 20 ng/mL Activin A (Peprotech), 20 ng/mL rhLIF (Peprotech), 0.5% KSR (Life Technologies), and 1 μM PD0325901 (Axon Medchem). After approximately 8-10 days, cells were detached using Accutase (Sigma-Aldrich), and subsequent centrifugation was performed in fibroblast medium composed of DMEM (Sigma-Aldrich) supplemented with 10% FBS (Corning), L-Glutamine (2mM, Sigma-Aldrich), 1% non-essential amino acids (Sigma-Aldrich), penicillin-streptomycin (Sigma-Aldrich), and 0.1 mM beta-mercaptoethanol (Sigma-Aldrich). The cells were then re-plated after passage through a 40μm cell strainer in 5i/LAF medium on irradiated CF-1 MEFs. The established naïve human ES cell lines were cultured on irradiated CF-1 MEFs at a density of 2.5×10^6^ cells per 9.5 cm² in 5i/LAF medium and were passaged every 6-7 days. The cells were provided with fresh medium daily. These naïve human ES cells were maintained under low oxygen conditions (5% O2) at 37°C.

### Three germ layer human ES differentiation

Human mesoderm progenitors were generated from human primed ES cells employing the STEMdiff™ Mesoderm differentiation Kit (Stem Cell Technologies), as per the manufacturer’s instructions.

For the derivation of human endoderm progenitors from human primed ES cells, the STEMdiff™ Definitive Endoderm Differentiation Kit (Stem Cell Technologies) was utilized following the manufacturer’s instructions.

To induce neural progenitors, human primed ES cells were cultivated in Neural Induction (NI) medium. This medium composition included a 1:1 ratio of Advanced DMEM/F12 medium and Neurobasal medium, supplemented with 1X N2 supplement (Life Technologies), 1X B27 supplement (Life Technologies), L-Glutamine (2 mM, Sigma-Aldrich), 4 μM CHIR99021 (Axon Medchem), 3 μM SB432542 (StemCell Technologies), 5 μg/mL BSA, 10 ng/mL human LIF (Peprotech), and 0.1 μM Compound E (γ-Secretase Inhibitor XXI, Peprotech). After a 9-day incubation period, the cells were passaged onto Matrigel-coated dishes using Accutase (Sigma-Aldrich). They were then maintained in the NI Medium, with the omission of Compound E. For the purpose of neuronal differentiation, the neural progenitors were cultured on Matrigel-coated dishes, utilizing DMEM/F12 medium (Sigma-Aldrich), 1X N2 supplement (Life Technologies), 1X B27 supplement (Life Technologies), 300 ng/mL cAMP (Sigma-Aldrich), and 0.2 μM vitamin C (Sigma-Aldrich).

### Endoderm progenitor maturation

hPSCs were sequentially differentiated towards anteriormost primitive streak, definitive endoderm, and then mature endoderm progenitors, as previously described^142^. Briefly, the following media compositions were used on each day of differentiation: Day 1: CDM2 base media supplemented with 100 ng/mL Activin A (Peprotech) + 3 μM CHIR99021 (Axon MedChem) + 20 ng/mL FGF2 (Peprotech) + 50 nM PI-103 (Tocris). Day 2: CDM2 base media supplemented with 100 ng/mL Activin A + 250 nM LDN-193189 + 50 nM PI-103. Day 3: CDM3 base media supplemented with 20 ng/mL FGF2 + 30 ng/mL BMP4 (R&D Systems) + 75 nM TTNPB (Selleck) + 1 μM A-83-01 (Reprocell). Day 4–6: CDM3 base media supplemented with 10 ng/mL Activin A + 30 ng/mL BMP4 + 1 μM Forskolin (Sigma-Aldrich).

The composition of CDM2 basal medium is: 50% IMDM + 50% F12 + L-Glutamine (2 mM, Sigma-Aldrich), penicillin-streptomycin (Sigma-Aldrich) + 1 mg/mL polyvinyl alcohol (Sigma) + 1% v/v chemically defined lipid concentrate (Thermo Fisher) + 450 μM 1-thioglycerol (Sigma-Aldrich) + 0.7 μg/mL recombinant human insulin (Sigma-Aldrich) + 15 μg/mL human transferrin (Sigma). The composition of CDM3 basal medium is: 45% IMDM + 45% F12 + L-Glutamine (2 mM, Sigma-Aldrich), penicillin-streptomycin (Sigma-Aldrich) + 10% KnockOut serum replacement (Thermo Fisher) + 1 mg/mL polyvinyl alcohol (Sigma-Aldrich) + 1% v/v chemically defined lipid concentrate (Thermo Fisher).

### Hepatocyte differentiation

For hepatic differentiation, endoderm cells were differentiated for 4 weeks in the SFD-based hepatic induction medium supplemented with ascorbic acid (50 μg/ml, Sigma), monothioglycerol (4.5×10^-4^ M, Sigma-Aldrich), BMP4 (50 ng/ml), bFGF (10 ng/ml), VEGF (10 ng/ml), EGF (10 ng/ml), TGFα (20 ng/ml, Peprotech), HGF (100 ng/ml, Peprotech), Dexamethasone (1×10^-7^ M, Sigma-Aldrich) and 1% DMSO (Sigma). SFD serum free medium consists of 75% IMDM (Life Technologies), 25% Ham’s F12 (Cellgro), 0.5× N2-Supplement (Gibco), 0.5× B27 without retinoic acid (Gibco), 0.1% BSA (Sigma-Aldrich), 50 μg/ml ascorbic acid phosphate magnesium (Wako), and 4.5 x 10^-4^ M monothioglycerol.

### PGCLCs differentiation

BTAG human iPSCs were maintained on iMatrix-511 (Reprocell) in Stemfit medium (Reprocell) supplemented with 100 ng/ml of bFGF (Peprotech). PGCLCs induction was achieved as previously described^130^. The iMeLCs were induced by plating 2.0X10^5^ hiPSCs per well of a 12-well plate coated with human plasma fibronectin (Millipore, FC010) in GK15 medium (GMEM (Life Technologies) supplemented with 15% KSR, 0.1 mM NEAA, 2 mM L-glutamine, 1 mM sodium pyruvate, and 0.1 mM 2-mercaptoethanol) containing 50 ng/ml of ActivinA (Peprotech), 3 μM CHIR99021, and 10 μM of ROCK inhibitor (Y-27632, Axon Medchem). The hPGCLCs were induced by plating 4X10^3^ iMeLCs per well of a cell repellent V-bottom 96-well plate (Greiner bio-one, 651970) in GK15 supplemented with 1,000 U/ml of hLIF (Millipore, #LIF1005), 200 ng/ml of BMP4 (Humanzyme), 100 ng/ml of SCF (R&D Systems, 455-MC), 50 ng/ml EGF (R&D Systems, 236-EG), and 10 μM of the ROCK inhibitor.

### HEK293T culture

HEK293T cells (human, female) were obtained from ATCC (Cat# CRL3216). These cells were maintained at a temperature of 37°C and were cultured in DMEM medium (Life Technologies), supplemented with 10% FBS (Corning), L-Glutamine (2 mM, Sigma-Aldrich), Penicillin (100 U/mL, Sigma-Aldrich), and Streptomycin (100 mg/mL, Sigma-Aldrich).

### Lentivirus production and transduction

HEK293T cells were co-transfected with a transfer plasmid along with packaging plasmids VSV-G and D8.9, using a calcium phosphate transfection method. Subsequently, viral supernatants were harvested within a time frame of 24 to 32 hours following transfection. These harvested viral supernatants underwent concentration through ultracentrifugation at a speed of 21,000×g for a duration of 2 hours, all performed at a temperature of 4°C. Once concentrated, the viruses were resuspended in Opti-MEM (Gibco). If not intended for immediate use, the concentrated viruses were stored at a temperature of -80°C.

### Generation of GFP-LSM14A cells

For HEK293T cells, mouse ES cells, and mouse embryonic fibroblasts, pEGFP-LSM14A from Weil lab^49^ was assembled into pLV-EF1a-Ires-Blast (Addgene # 85133).

For human ES cells, pEGFP-LSM14A was assembled into pAAVS1-tet-iCas9-BFP2 (Addgene #125519), replacing iCas9-BFP2 sequence with pEGFP-LSM14A.

### Generation of DDX6 degron mouse ES cells

To introduce a sequence encoding FKBP12F36V-HA-2A-mCherry in place of the endogenous *Ddx6* stop codon, a first donor plasmid was created by integrating two 200 bp homology arms specific to the *Ddx6* gene into the pNQL004-SOX2-FKBPV-HA2-P2A-mCherry targeting construct (Addgene #175552). In conjunction with this donor plasmid, a *Ddx6*-targeting sgRNA (AGGTACATACGTGCTTGTTA) was cloned into the pSpCas9 (BB)-2A-Hygro plasmid (Addgene #127763). Both donor plasmids were transfected into GFP-LSM14A mouse ES cells using the Lipofectamine 3000 transfection method. Cells exhibiting stable mCherry expression were subsequently isolated using fluorescence-activated cell sorting (FACS) to establish single-cell clones. Homozygous insertion was confirmed by genotyping PCR. For this purpose, a forward primer (GCTGGGGACAGAGATCAAACC) and a reverse primer (GTAGGGCTATGCGGCCCTAC) within the DDX6 homology arms were utilized, along with a reverse primer specific to the mCherry sequence (ATCTGGGCAACCCCTTCTTCC). Loss of endogenous DDX6 upon dTAG-13 treatment was confirmed through western blot analysis.

### Generation of AGO2 KO mouse ES cells

The Ago2_KO cell line was established via a paired CRISPR/CAS9 approach implemented on wild-type (WT) mouse ES cells, as previously described by^115^.

Briefly, v6.5 mouse ES cells underwent transfection with pX458-sgRNA_Ago2_3/4 plasmids (Addgene # 73531 and 73532). GFP-positive cells were isolated through single-cell sorting in 96-well cell plates. To verify the deletion, PCR using the Ago2KOΔEx1_Fw (GAAGGCGAAAAAGCCTCCCC) and Ago2KOΔEx1_Rev (GAGCTAGCTTCCCGTCGC) primers was conducted. Positive clones were expanded and verified by western Blot and RT-qRT-PCR analyses.

### shRNA-mediated miRNA gene silencing

The pLKO-sh*DDX6* (TRCN0000074696) vector, designed to target the human *DDX6* gene, was sourced from the Molecular Profiling Laboratory of the MGH Cancer Center.

To target the mouse Ddx6 gene, the shERWOOD UltramiR Lentiviral shRNA was procured from Transomic Technologies, as previously described^48^.

To accomplish *Nudt21* gene silencing, oligonucleotide pairs encoding shRNA sequences specific to *Nudt21* (GGACAACTTTCTTCAAATT) were annealed and cloned into the pSICOR-mCherry-puro vector (Addgene #31845). The efficacy of knockdown was validated by qRT-PCR.

In the context of miRNA knockdown experiments, miR-300 and control miRNA inhibitors (miRNA Power Inhibitors, Qiagen) were directly introduced into the cell culture media. These inhibitors were added at a final concentration of 50nM, as per the manufacturer’s recommendations.

### Generation of TPRX1-GFP human ES cells

For knock-in TPRX1–EGFP generation, sgRNA (GCTCCCGAGCTAGTTTGGCG) targeting the insertion site of the donor construct containing homologous arms flanked by an EGFP–puromycin cassette, kindly gifted by the Esteban lab^83^, was cloned into pDonor plasmid. Donor plasmids were electroporated into UCLA-4 human ES cells using the Neon Transfection System.

### Generation of miRNA reporter cell lines

Plasmid pNZ176, containing Halo-NLS-MCP, was a kind gift from Tim Stasevich. Halo-NLS-MCP fragment was PCR amplified from this vector in parallel with PCR amplification of Ires BFP fragment from TRE KRAB Cas9 Ires BFP plasmid (Addgene # 85449) using Phusion polymerase (New England Biolabs) according to the manufacturer’s recommendations. The resulting fragments were used for NEBuilder® HiFi DNA Assembly (New England Biolabs). Final vector was transfected along with PBase vector into Nanog KO^143^ mouse ES cells using the Lipofectamine 3000 transfection method. BFP-positive cells were isolated using fluorescence-activated cell sorting (FACS).

MS2 reporter construct for *Nanog* was generated using a multistep cloning approach. First, *Nanog* was PCR amplified from the pMXs-Nanog (Addgene #13354)^144^ plasmid using Phusion polymerase (New England Biolabs) according to the manufacturer’s recommendations. Primers to amplify *Nanog* were CGCTGTGATCGTCACTTGGCGCCGCCATGAGTGTGGGTC and CGCTGTGATCGTCACTTGGCCCACCATGAGTGTGGGTC. PCR product was treated with DpnI (New England Biolabs) and purified using a PCR cleanup kit (Qiagen). The resulting *Nanog* fragment was used for Gibson assembly (New England Biolabs) with pUB_smFLAG_ActB_MS2 (Addgene #81083)^145^ digested with NotI and NheI (New England Biolabs). The resulting pNANOG-MS2 plasmid was used for Gibson assembly (New England Biolabs) with the Let-7^WT^ and Let-7^MUT^ sequences amplified from the RL-6×B and RL-6×BMUT vectors (a kind gift of Marc R. Fabian)^121^. Finally, the NANOG-Let-7^WT^-MS2 and NANOG-MS2-Let-7^Mut^-MS2 fragments were cloned into the PiggyBac vector PB-TRE-EGFP-EF1a-rtTA-Blasti digested with Nhe1 and Kpn1. Final vectors were used to transfect Nanog KO HALO-NLS-MCP IRES BFP.

### Western blot

Cells were lysed in RIPA buffer (50 mM Tris-HCl (pH 8.0), 150 mM NaCl, 0.1% SDS, 0.5% sodium deoxycholate, 1% Triton-X-100, 1 mM EDTA, protease inhibitors, and benzonase). Lysates were subjected to standard western blotting using the following antibodies: rabbit anti-human/mouse DDX6 (1:5000, Novus Biologicals), Rabbit anti-Argonaute-2 (1:1000, C34C6; Cell Signaling Technologies), HRP-rabbit anti-human/mouse β-Actin (1:3000, Cell Signaling), Goat, anti-rabbit-HRP-conjugated (Thermo Fisher Scientific, 1:2,000 dilution), Rabbit, anti-mouse-HRP-conjugated (Thermo Fisher Scientific, 1:2,000 dilution).

### Immunocytochemistry and confocal microscopy

Cells were fixed using 4% methanol-free formaldehyde for a duration of 15 minutes at room temperature. After fixation, the cells were permeabilized and simultaneously blocked using a solution of 0.1% Triton-X and 10% donkey serum. Subsequently, primary antibodies were applied to the cells and allowed to incubate at 4°C overnight. Following this step, the cells were stained with Alexa Fluor-conjugated secondary antibodies, at room temperature for a duration of one hour. Where mentioned, Phalloidin dye (Abcam) was combined with the secondary antibody to facilitate the labeling of membranes. Nuclei were counterstained using DAPI. The slides were mounted using ProLong Diamond (Life Technologies) and subsequently allowed to cure at room temperature. The following primary antibodies were used in this study: EDC4 (1:50, Abcam), GFP (1:400, Aves Lab), KLF17 (1:100, Sigma), NANOG (D73G4) (1:300, Cell Signaling), BRACHYURY (T) (1:50, Cell Signaling), FOXA2 (1:400, Cell Signaling), SOX1 (1:200, R&D Systems), βIII-TUBULIN (TUJ1) (1:200, Biolegend), METTL3 (1:100, Proteintech). Imaging was carried out using confocal microscopy equipped with either a 40× or 63× objective lens on an LSM 900 with Airyscan 2 (Zeiss). The acquired images were then subjected to analysis using ImageJ software.

P-bodies were counted manually to calculate number of RNP particles per cell.

### Immunofluorescence and Fluorescence in situ hybridization (IF-FISH)

Cells were fixed using 4% methanol-free formaldehyde for a period of 30 minutes on ice. Subsequently, cells underwent permeabilization through a 0.1% Triton-X100 solution for 30 minutes at room temperature. For antibody labeling, cells were immunolabeled with chicken anti-GFP (1:400, Aves Lab) at 37°C for 1 hour in 2×SSC buffer, followed with Alexa Fluor 488-conjugated donkey anti-chicken IgG (H+L) (1:500, Life Technologies) for a duration of 30 minutes at room temperature. Afterward, a post-fixation step was carried out using 4% methanol-free formaldehyde for 15 minutes at room temperature. Following this, the cells were labeled using a hybridization buffer containing POLK (SMF-1063-5 Stellaris, LGC Biosearch Technologies) or control probes (T30-QUASAR 570-1 Stellaris) at 37°C overnight.

After washes, nuclei were counterstained using DAPI, and samples were imaged within the next three days. The imaging was conducted using a Cytivia DVLive system with an Olympus PlanApo N 60×/1.42NA oil objective, capturing 6 fields per well and collecting a total z-stack of approximately 15μm with optical spacing of 0.25µm. Images were then deconvolved using a conservative restorative algorithm, and then 16-bit TIFF files were used for image quantification. Quantification was performed using FISH-quant^146^.

### RNA Dot blot

200ng of RNA was loaded on a positively charged nylon transfer membrane (GE Healthcare). Membrane was UV cross-linked at 1200 J/m2, and subsequently blocked with gentle agitation for 1 hr at room temperature. Membrane was incubated with m6A primary antibody (Sigma, 1:3000) with gentle agitation overnight at 4°C. Membrane was then incubate with HRP-conjugated secondary antibody (Anti-rabbit IgG, HRP-linked Antibody) at 1:3000 with gentle agitation for 1 hr at room temperature. Methylene blue staining was used to normalized sample loading.

### Colony number quantification

ImageJ^147^ was used to quantify colonies based on alkaline phosphatase (AP) images as previously described^140^. A region of interest of identical size was selected in each well using the oval tool. The “find edges” command was executed and the “color threshold tool” was used with the following parameters: “default thresholding method”, “red” as the threshold color, “HSB” as the color space, and “dark background.” “Analyze particles” was used to quantify the number of AP-positive colonies and the area of those colonies using the following parameters: “Size (pixel^2)=9-infinity”; “Circularity=0–1”; “Exclude on edges” was applied; “Include holes” was applied. The settings were identical for all samples processed.

### Quantitative RT-PCR (qRT-PCR)

RNA extraction from cells was performed using the Monarch Total RNA Miniprep Kit (New England Biolabs). After RNA extraction, reverse transcription to cDNA was performed utilizing the LunaScript RT SuperMix Kit (New England Biolabs), according to the manufacturer’s instructions. For qRT-PCR analysis, a master mix was prepared comprising Luna Universal qPCR Master Mix (New England Biolabs) along with pre-designed primers (Sigma-Aldrich) at a concentration of 0.5μM for the target genes. The qPCR reactions were conducted on a CFX96 Real-Time PCR Detection System (Bio-Rad). The cycling conditions for the qRT-PCR reactions were as follows: an initial denaturation step at 95°C for 1 minute, followed by 40 cycles of denaturation at 95°C for 15 seconds, and annealing/extension at 60°C for 30 seconds. Primers are available upon request.

### P-body purification

P-body purification was performed as previously described. Briefly, GFP-LSM14A cells were subjected to lysis for a duration of 20 minutes on ice using lysis buffer (comprising 50 mM Tris at pH 7.4, 1 mM EDTA, 150 mM NaCl, and 0.2% Triton X-100), supplemented with 65 U/mL of the ribonuclease inhibitor RNaseOut (Promega) and an EDTA-free protease inhibitor cocktail (Roche Diagnostics). Following this step, the lysates underwent centrifugation at 200×g for 5 minutes at 4°C to remove nuclei from the mixture. Residual DNA was removed by incubation in the presence of 10 mM MgSO4, 1 mM CaCl2, and 4 U/mL of RQ1 DNase (Promega) for 30 min at room temperature. Following centrifugation at 10000×g for 7 min at 4°C, pellets were resuspended in 40μL of lysis buffer containing 80 U of RNaseOut (Promega) to generate the cytoplasmic fraction. P-bodies were then isolated from this cytoplasmic fraction using FACSAria-based sorting. Following sorting, the P-body samples were pelleted via centrifugation at 10,000×g for 7 minutes at 4°C. These pellets, alongside aliquots of the corresponding pre-sort cytoplasmic fraction, were stored at a temperature of -80°C.

### RNA-seq

For P-body-seq, RNA was isolated from P-body and cytoplasmic fractions using the miRNeasy Micro Kit (QIAGEN). cDNA libraries were constructed using the SMART-Seq V4 ultra + Nextera XT kit for Illumina (Takara Bio). Where indicated in the text, libraries were prepared using SnapTotal-Seq protocol as previously described^66^. Libraries were sequenced with paired-end 150-bp reads.

### Small RNA-seq

Purified RNA from P-body and unsorted fractions of naïve and primed mouse ES cells, primed human ES cells, and human neural progenitors was mixed with RNA loading dye (New England Biolabs; B0363S) by equal volume, and incubated at 75 °C for 5 min. The mixture was loaded into 15% (wt/vol) urea polyacrylamide gel and ran in a 1× TBE running buffer at 200 V until the bromophenol blue reached the bottom of the gel. After staining with SYBR Gold solution (Invitrogen; S11494), the gel that contained 15-50 nucleotides RNAs was excised based on small RNA ladders (New England Biolabs) and Takara (3416) and eluted in 0.3 M sodium acetate (Invitrogen) and 100 U ml^-1^ RNase inhibitor (New England Biolabs) overnight at 4 °C. The sample was then centrifuged for 10 min at 12,000g (4 °C). The aqueous phase was mixed with pure ethanol, 3 M sodium acetate and linear acrylamide (Invitrogen) at a ratio of 3:9:0.3:0.01. Then, the sample was incubated at -20 °C for 2 hours and centrifuged for 25 minutes at 12,000g (4 °C). After removing the supernatant, the precipitation was resuspended in nuclease-free water, quantified, and stored at -80°C or used for further processing. The size-isolated RNAs were incubated in 50μL reaction mixture containing 5 μL 10× PNK buffer (New England Biolabs), 1 mM ATP (New England Biolabs), 10 U T4PNK (New England Biolabs) and RNA at 37 °C for 20 min. Then, the mixture was added to 500 μL TRIzol reagent to perform the RNA isolation procedure. After purification, the RNAs were incubated in 50 μl reaction mixture containing 50 mM HEPES (pH 8.0), 75 μM ferrous ammonium sulfate (pH 5.0), 1 mM α-ketoglutaric acid, 2 mM sodium ascorbate, 50 mg l^-1^ bovine serum albumin (Sigma-Aldrich; A7906-500G), 2,000 U ml^-1^ RNase inhibitor, and equal concentrations of AlkB and RNA at 37°C for 30 min. Then, the mixture was added to 500μL TRIzol reagent to perform the RNA isolation. Small RNA libraries were constructed using the NEBNext Small RNA Library Prep Set for Illumina (New England Biolabs). Libraries were amplified and sequenced using the SE100 strategy on the Illumina system by the University of California, San Diego IGM Genomics Center.

### Polysome profiling

Sucrose gradients were made by preparing 10% and 50% (w/w) sucrose solutions in polysome buffer (15 mM Tris-HCl, 15 mM MgCl2, 0.3M NaCl, ph 7.4) supplemented with 100ug/mL Cycloheximide (CHX, Tocris). 5.9mL 50% sucrose was loaded into each centrifuge tube (Polyallomer 14×95mm, Beckman #331372), followed by 10% sucrose on top, then placed in a gradient maker (Gradient Master, Biocomp model 108), with the program set to: SHORT SUCR 10–50% WW 1st (Time 1:55; Angle 81.5, Speed 25 rpm; 1 cycle). Prepared gradients were stored at 4°C for 2 hours prior to use.

DDX6 degron 2i mouse ES cells (2× 10cm plates for each biological replicate) were grown until confluent. To induce degron, dTag13 (dTAG13 Sigma) was added to a final concentration of 1μM. After 6 hours, cells were treated with 100 μg/ml CHX for 10 minutes at 37°C. Cells were washed once with 1X PBS supplemented with 100 μg/mL CHX, and harvested using Accutase supplemented with 100 μg/mL CHX. After quenching, cells were spun down at 200g for 5 min, and washed two times with 10 mL 1X PBS supplemented with 100 μg/mL CHX. Cell pellets were resuspended in 400 μL lysis buffer (polysome buffer with 1% Triton-x 100, 0.1 U/μL RNase OUT (Invitrogen), 0.125 U/μL DNase I (Zymo Research), incubated on ice for 10 minutes, and then centrifuged at 12,000×g at 4°C for 10 min. 50 μL of each lysate was kept for total RNA samples, and the remaining 350 μL were used for fractionation. Control cells were harvested in the same manner alongside dTag13 treated cells.

For fractionation, 350 μL of the lysate was loaded onto each gradient and centrifuged at 30,000 RPM for 2hrs at 4°C (Beckman Optima-XE 90, SW41 rotor). 300 μL fractions were collected from the top using a fractionator with a UV monitor (Density Gradient Fractionation System—Teledyne ISCO). Ribosome-containing fractions were identified by the UV spectrum, and pooled to make up the ribosome-bound pool. RNA from all samples was extracted using the Direct-zol RNA Miniprep kit (Zymo Research, R2050). Prior to loading on the columns, Trizol-LS (Invitrogen) was added to sucrose at 1:1 volume, followed by 1:1 100% ethanol. RNA integrity and concentration were checked using Agilent tape station, and libraries were prepped from a starting input of 500ng using the NEB Ultra II Directional RNA Library Prep Kit for Illumina (NEB). Libraries were quantified using the NEBNext Library Quant Kit for Illumina (NEB) and pooled to equal amounts of each library. Final libraries were sent to Novogene for sequencing and were sequenced at a depth of >35M reads.

### RNA-seq analysis

RNA-seq was analyzed by mapping reads using hisat2.1.0^148^ to the mouse mm10 genome assembly, human hg38 assembly, or the chicken gal6 assembly, all retrieved from UCSC. Gene counts were annotated using featureCounts v1.6.2 using the corresponding UCSC refGene annotation.

To determine P-body enriched and depleted genes, lowly expressed genes (<10 reads over all samples) were filtered out. Then, DESeq2 v1.40.2 was used to calculate differential expression between the P-body and cytoplasmic fractions, setting an FDR of p<0.05 as a threshold for significance. P-body to cytoplasm ratios are represented as log2FC (p-body/cytoplasm), where a positive log2FC indicates p-body enrichment, and a negative log2FC indicates p-body depletion (cytoplasm enriched).

Gene set enrichment analyses were performed using fgsea v1.26.0 R package with nperm=1000, with the following gene sets (human neurons^149^, human naïve^2^, mouse 2C^5^, human 8C^3^, human 8CLC^3^, ZGA^3^, PGCLCs^130^). Gene ranks were determined by the stat parameter in the results tables generated by DESeq2. Gene ontology analysis was performed using enrichGO, from the clusterprofiler package. Top GO terms were determined by padj value. Orthologs were determined using Ensembl Biomart genes v.108. MiRNA targets were obtained from the miRDB database^7, 150^, where targets with a threshold score greater than the 75^th^ percentile was deemed a target. To infer targets of RNA binding proteins, the POSTAR3/CLIPDb database was used. PCA plots were generated using the prcomp function in R on all genes, scaled. Venn diagrams were made using the euler function from the eulerr R package (version 7.0.2).

SnapTotal-Seq data were processed using the nf-core RNA-seq pipeline (version 3.9) within a Singularity container. Reads were aligned to the mouse mm10 genome assembly and the human hg38 genome assembly, both of which were retrieved from UCSC with corresponding UCSC refGene annotations. Raw counts were obtained from Salmon (v1.10.2), and the resulting DDS objects from nf-core were exported to R for downstream analysis. For differential gene expression analysis, genes with fewer than 10 reads across all samples were filtered out, and DESeq2 was used to assess differential expression between P-body and cytoplasmic fractions, applying an FDR threshold of p<0.05 for significance. In human HEK293T and mouse naïve ES cells, genes significantly enriched in P-bodies were compared to those enriched in P-bodies in SMART-Seq data, and overlaps were visualized with the ggVenn package in R. For GSEA of 2C genes in naïve mouse ES cells, the fgsea package (v1.26.0) in R was used with a predefined mouse 2C gene set, with gene ranks based on the stat parameter from DESeq2 results.

Gene coverage plots were generated using PyCoverage (geneBody_coverage.py from the RSeQC package). First, BED12 files were created for P-body-enriched genes, cytoplasm-enriched genes, and all transcripts for each cell type (gene sets defined previously in RNA-seq methods). Gene annotation files in BED format were downloaded from the UCSC Genome Browser (https://genome.ucsc.edu/) for mm10 and hg38 genome assemblies, selecting Genes and Gene Predictions group, NCBI RefSeq track, and the UCSC RefSeq (refGene) table. These files were converted to BED12 format with custom R code, where block sizes were calculated as the difference between exon start and end positions, and block starts were calculated as the difference between exon start and transcript start. Only the longest isoform per gene was retained. Filtered genome BED12 files for the specified gene lists were then used in the geneBody_coverage.py command, with BAM files for each fraction (two replicates each) analyzed per cell type and species. Resulting outputs were plotted in R, displaying read counts across gene bodies.

To generate specific gene sets for cell type comparisons lacking publicly available data, we performed DESeq2 analysis on the sequencing results from the cytoplasmic fractions of each cell type. Gene sets were generated in human samples for the following comparisons: primed vs. endoderm progenitor, primed vs. mesoderm progenitor, naïve vs. primed, and primed vs. mature endoderm progenitors. Each gene set included genes with a log2 fold change greater than 2.5 and an adjusted p-value (padj) <0.05 for each comparison.

To calculate GC content and transcript length distributions for P-body- and cytoplasm-enriched genes, we used the biomaRt package to access cDNA sequences from Ensembl (hsapiens_gene_ensembl). Sequences were retrieved using the *cdna* attribute, with GC content calculated as the percentage of G and C nucleotides and transcript length as sequence length in nucleotides. For uniformity, only the longest isoform per gene was retained. Density plots for GC content and transcript length were generated using ggplot2, and two-sample t-tests were conducted to assess statistical significance.

### Proteomics analysis

Cell pellets were resuspended in lysis buffer (8 M urea, 150 mM NaCl, 100 mM Tris pH 8) and sonicated for 20s at 20% amplitude. Samples were incubated with dithiothreitol (DTT) to a concentration of 5 mM for 30 minutes at room temperature. Iodoacetamide (IAA) was added to a final concentration of 10 mM, and samples were incubated for 30 minutes in the dark. A final incubation with 10 mM DTT was performed for 30 minutes at room temperature. An automated SP3^151^ digestion protocol was performed on the King Fisher Flex (Thermo). Briefly, 20 uL of MagReSyn Amine beads per sample were washed twice with 60 uL of HPLC water. After washing, beads were resuspended in 20 uL HPLC water. Protein lysates were added to a deep well plate, where they were diluted to 100 uL with water. 20 uL of beads were added to each well, and each sample was diluted with 100% acetonitrile (ACN) to a final ACN concentration of 70% to induce aggregation of proteins to the beads. The beads were then washed thrice with 150uL of 95% ACN, followed by two washes with 150uL of 70% ethanol. Finally, beads were deposited in a plate containing trypsin (Promega, 1:100 enzyme:protein ratio) and lysC (Wako, (1:100 enzyme:protein ratio) in 50 mM ammonium bicarbonate. Beads were incubated overnight at 37°C with 800 rpm mixing and the resulting supernatant, containing the digested peptides, was transferred to a PCR filter plate to remove residual beads. After filtration, protease activity was quenched by acidification with formic acid to a final volume percentage of 1%. This first elution was then dried by vacuum centrifugation. The elution of peptides off the beads was repeated by the addition 50 uL of water to the beads with shaking for 1h at RT. Following shaking, the liquid was collected from the beads, filtered, and combined with the first elution, and again dried by vaccum centrifugation.

The resulting peptides were separated on a PepSep reverse-phase C18 column (1.9 μm particles, 15 cm, 150 mm ID) (Bruker) with a gradient of 3–28% buffer B (0.1% formic acid in acetonitrile) over buffer A (0.1% formic acid in water) over 67 minutes, an increase to 40% B in 5 minutes, and held at 95% B for 8 minutes. Eluting peptide cations were analyzed by electrospray ionization on an Orbitrap Eclipse (Thermo Fisher Scientific). For data-independent acquisition (DIA) analysis, MS1 scans of peptide precursors were performed at 120,000 resolution (200 m/z) over a scan range of 350-1050 m/z, with an AGC target of 250% and a max injection time of 100 ms. MS2 scans were collected over 350-950 m/z in 15 m/z isolation windows with a 0.5 m/z overlap. Maximum injection time was set to auto, with AGC set to standard, and an MS2 resolution of 15,000. Higher energy collisional dissociation (HCD) was performed at a NCE of 28%. Six gas-phase fractionation (GPF)^152^ -DIA runs were collected from a pooled sample of all conditions. tSIM MS1 scans were performed at 120,000. tMS2 scans were collected at 30,000 fragment resolution, an AGC target of 1e6, a maximum ion injection time of 60 ms, and a NCE of 26. Data was collected using 4*m*/*z* precursor isolation windows in a staggered-window pattern, with each GPF fraction covering an approximately 100 m/z range.

All data was searched against the Uniprot mouse database (downloaded 06/11/24) using the GPF library in Spectronaut (Biognosys, version 19.0). Default settings, including trypsin digestion, variable modifications of methionine oxidation and N-termini acetylation, and fixed modification of cysteine carbamidomethylation, were used. DIA runs were filtered to obtain a false discovery rate of 1% at the peptide spectrum match and protein level^153^. Quantitative analysis was performed in the R statistical programming language using the artMS package (release 3.20) and MSstats (version 3.20)^154^. For MSstats analysis, normalization_method was set to equalizeMedians, and censored missing values were imputed by the Accelerated Failure Time model.

### Alternative Polyadenylation analysis

Alternative polyadenylation was analyzed using LABRAT (v1.0)^155^. Paired-end FASTQ files were input to LABRAT’s runSalmon mode to quantify transcripts, using the reference genome FASTA (GRCh38) and GFF annotation (gencode v28) files with the lasttwoexons option specified. The calculatepsi mode was subsequently run on the Salmon output to compare alternative polyadenylation usage between cytoplasmic and P-body fractions.

### SLAM-Seq analysis

SLAM-Seq analysis was conducted to assess the correlation between P-body enrichment and mRNA half-life, using publicly available SLAM-Seq data from HEK293T cells^156^, mouse embryonic fibroblasts^157^, and mouse ES cells^90^. Additionally, poly(A) tail length, as an indicator of mRNA stability, was evaluated by comparing PAISO-Seq data from mouse ES cells^91^. For each dataset, a Pearson correlation was calculated.

### Motif Enrichment Analysis

To identify sequence motifs enriched in P-body transcripts, we used the STREME^158^ function from the MEME suite^159^ (version 5.5.7). For target sequences, FASTA files were generated containing the longest 3’ UTR of the top 100 P-body-enriched genes for each cell type, while control sequences consisted of FASTA files with the longest 3’ UTR of the top 100 P-body-depleted genes. To create these FASTA files, BED files were downloaded from the UCSC table browser for mm10 and hg38 annotations by selecting the Genes and Gene Predictions group, NCBI RefSeq track, and UCSC RefSeq (refGene) table. For extraction of 3’ UTR sequences only, the setting “create one BED record per 3’ UTR Exons” was applied, and the longest annotated 3’ UTR per gene was retained. These BED files were filtered to include only the top 100 P-body-enriched and top 100 P-body-depleted genes and then converted to FASTA format using the bedtools (v2.28.0) getfastafunction. Tomtom^159^ (MEME Suite) was subsequently run using the streme.txt output from STREME and the following target motif databases: RNA (DNA-encoded) Ray 2013 RNA Binding Protein dataset and miRBase^160^ (v22) Single Species miRNA (DNA-encoded) for the respective species.

### Small RNA-seq analysis

The raw reads were trimmed and annotated by performing SPORTS1.1.2 (parameters: sports.pl-M 1 -a -x GUUCAGAGUUCUACAGUCCGACGAUC -y AAGATCGGAAGAGCACACGTCT -l 15-L 45 -s) with the pre-compiled mouse annotation database (mm39 (for mouse), hg38 (for human)). Pairwise comparison of differentially expressed sncRNAs among different groups was performed using the R package DEseq2 with a threshold of FC > 2 and adjusted p<0.05.

### Polysome profiling analysis

RNA-seq from polysome containing samples and total RNA samples was analyzed using the nf-core/rnaseq pipeline (v 3.9), with default settings. Reads were aligned to the mouse mm10 genome assembly, retrieved from UCSC using the corresponding UCSC refGene annotation. Raw counts were obtained from salmon v1.10.2.

To determine differentially expressed genes in DDX6 knockdown samples, lowly expressed genes (<10 reads over all samples) were filtered out. Then, DESeq2 was used to call differential expression between control and dTag13-treated total RNA samples. An FDR of p<0.05 was used as the threshold for significance in differentially expressed genes.

For polysome fractionation analysis, genes were filtered to protein-coding genes using the mm10 refGene annotation. Ribosome occupancy) was calculated as the log2 (ribosome bound/total RNA), using DESeq2. Change in ribosome occupancy was calculated as ribosome occupancy degron-ribosome occupancy control. GSEA was performed as above using mouse gene sets from (2C)^5^, (Naïve)^6^, and (Primed)^7^.

### Translation efficiency analysis

Ribo-seq data was obtained from^1^. To download the Ribo-seq raw data, GSE133794 was downloaded from Gene Expression Omnibus (GEO) and processed using the nf-core/rnaseq pipeline (v 3.9). Reads were aligned to the mouse mm10 genome assembly, retrieved from UCSC using the corresponding UCSC refGene annotation. Raw reads were obtained from salmon v1.10.2. Reads were normalized by RPKM, and lowly expressed genes (<10 reads over all samples) were filtered out. Translation Efficiency (TE) was calculated as log2 (reads in ribo-seq/reads in RNA-seq) for each gene.

### Statistical Analysis

Quantitative data are presented as mean ± s.d. Statistical analyses were performed using Prism 9.3.0 software (GraphPad). Details for statistical analyses, including replicate numbers, are provided in the figure legends.

Bioinformatics analyses and statistics were done using R 4.3.1. Boxplots were made using ggplot2 version 3.5.1 with the ggpubr 0.6.0 extension, and show mean plus quartiles, and whiskers show minimum and maximum quartiles excluding outliers.

---

**Extended Data Figure 1.**
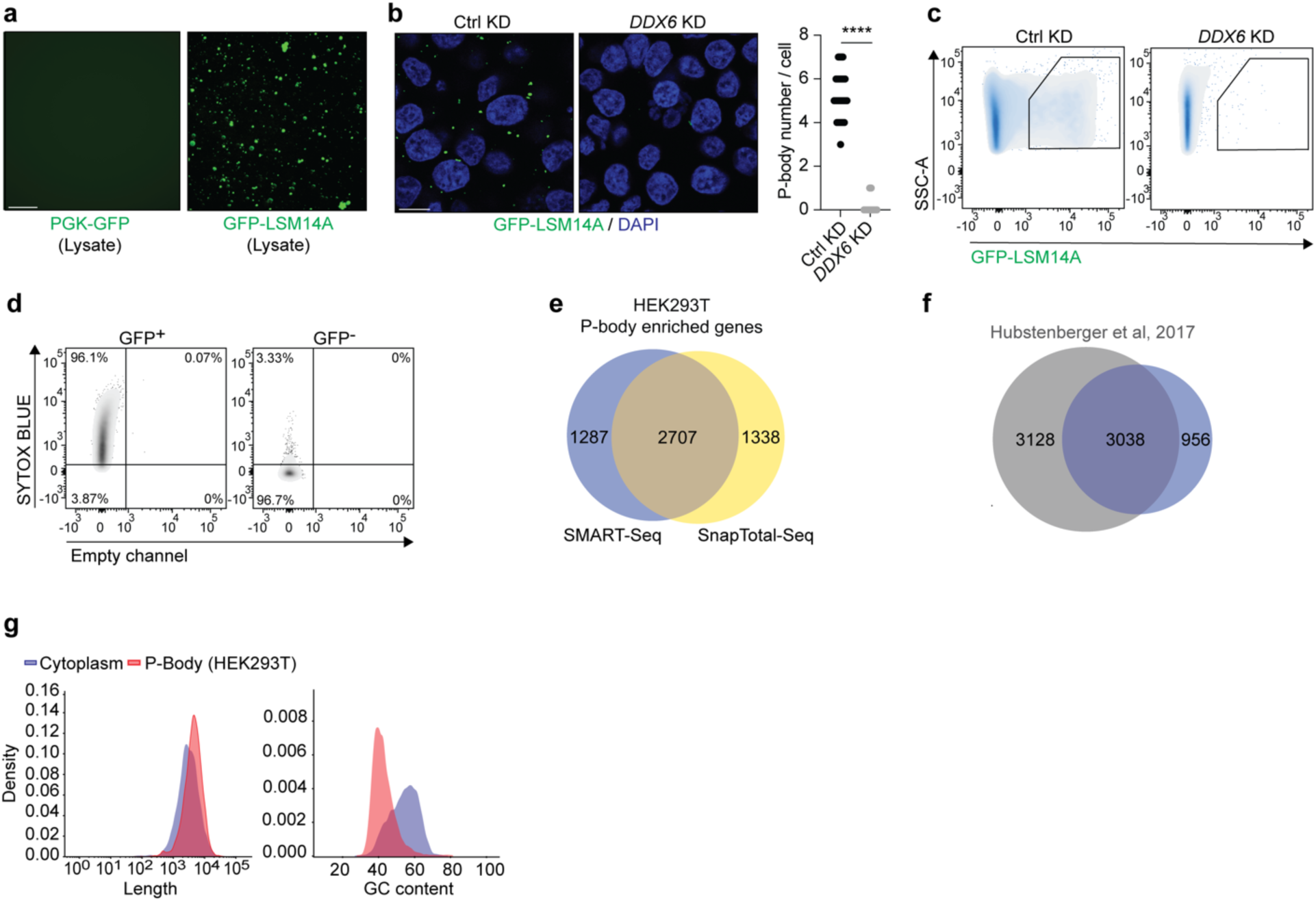
DDX6 suppression abolishes P-body formation. **(a)** Representative image of pre-sorted cell lysate containing GFP-LSM14A^+^ P-bodies (scale: 10μm). **(b)** Representative imaging of GFP-LSM14A puncta (green) in control and *DDX6* KD HEK293T cells. Nuclei were counterstained with DAPI (blue) (scale: 10μm) (left panel). P-body number in control (n=50 cells) and *DDX6* KD (n=50 cells) HEK293T cells (right panel). Unpaired Student’s t-test, mean ± s.d., ****: p<0.0001. **(c)** Representative flow cytometry plots showing gating for GFP-LSM14A^+^ P-bodies in control and DDX6 KD HEK293T cells. **(d)** Representative flow cytometry plots showing gating for SYTOX BLUE^+^ events in GFP^+^ and GFP^-^ gates of pre-sorted cytoplasmic fraction from HEK293T cells. **(e)** Venn diagram showing the overlap between P-body-associated mRNAs in HEK293T from SMART-Seq and SnapTotal-Seq. **(f)** Venn diagram showing the overlap between P-body-associated mRNAs in HEK293T of this study (blue) and a published dataset from^49^ (grey). **(g)** Length (left panel) and GC content (right panel) density plots of mRNAs in purified P-body and cytoplasmic fractions of HEK293T cells.

**Extended Data Figure 2.**
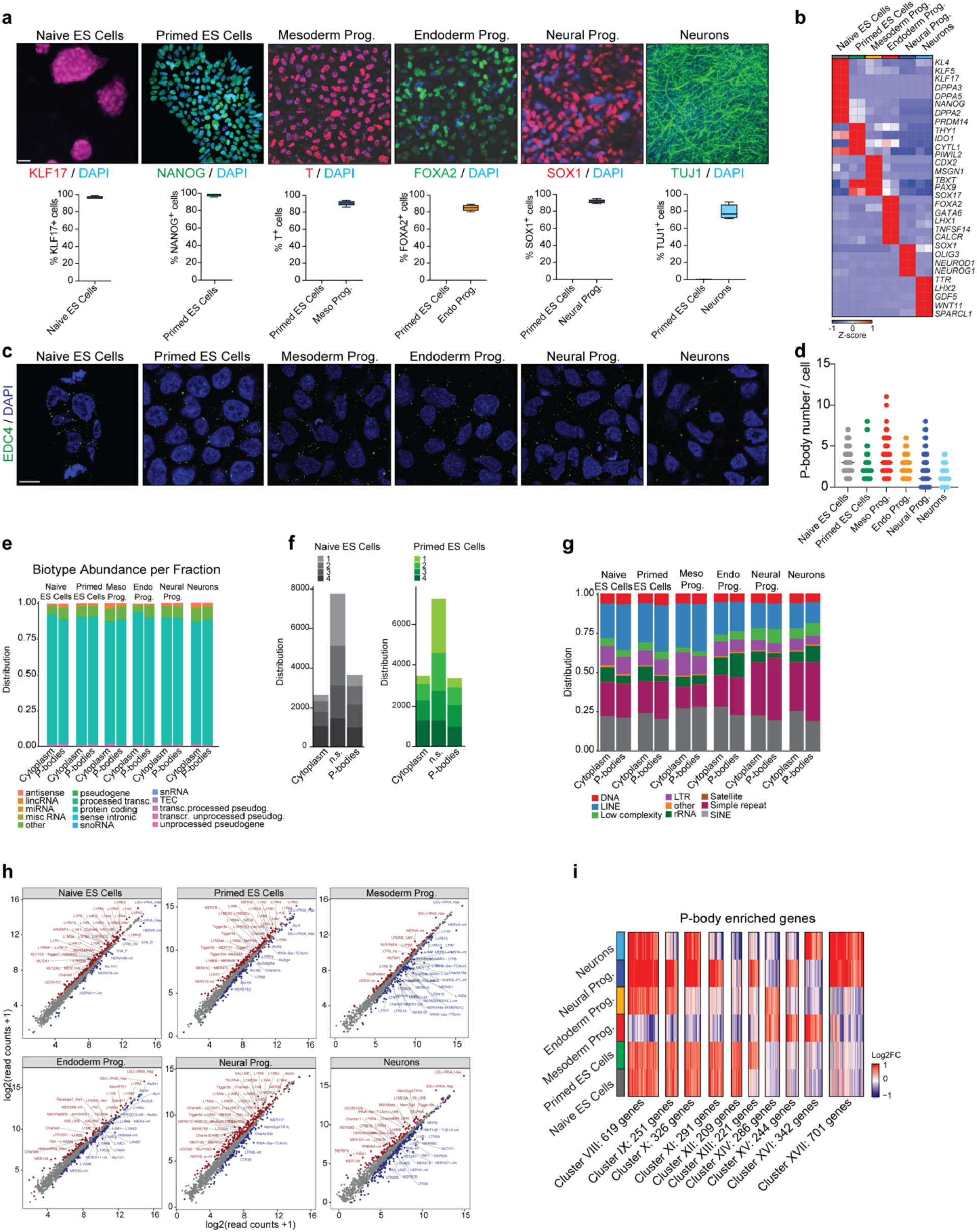
Directed differentiation of human ES cells permits analysis of P-body contents. **(a)** Representative IF images of lineage-specific markers for the indicated samples. Nuclei were counterstained with DAPI (blue) (scale: 25μm) (upper panel). Quantification of lineage-specific positive cells in the indicated samples (bottom panel). **(b)** Heatmap showing expression levels of lineage-specific genes for the indicated samples. (n=2). **(c)** Representative IF imaging of EDC4 puncta (green) in the indicated samples. Nuclei were counterstained with DAPI (blue) (scale: 10μm). **(d)** P-body number in the indicated samples. Naïve human ES cells (n=70 cells), primed human ES cells (n=90 cells), mesoderm progenitors (n=90 cells), endoderm progenitors (n=90 cells), neural progenitors (n=90 cells), neurons (n=90 cells). **(e)** Distribution of biotypes within the P-bodies and cytoplasm of each cell type. **(f)** Distribution of P-body enriched, cytoplasm enriched, and NS genes within each expression quartile. Quartiles are calculated by taking the distribution of counts per gene, so that each quartile has an equal number of genes, with Q1 being lowly expressed and Q4 being highly expressed. **(g)** Ratio of repetitive elements in purified P-body vs. cytoplasmic fractions of the indicated samples. **(h)** Differential expression analysis of transposable element expression in purified P-body vs. cytoplasmic fractions of the indicated samples. *red: p-adj<0.05, log2FC>0, blue: p-adj<0.05, log2FC<0. **(i)** Heatmap showing expression levels of differentially enriched mRNAs between purified P-body fractions of the indicated samples. Gene number in each cluster is indicated in the figure (n=2, p < 0.05).

**Extended Data Figure 3.**
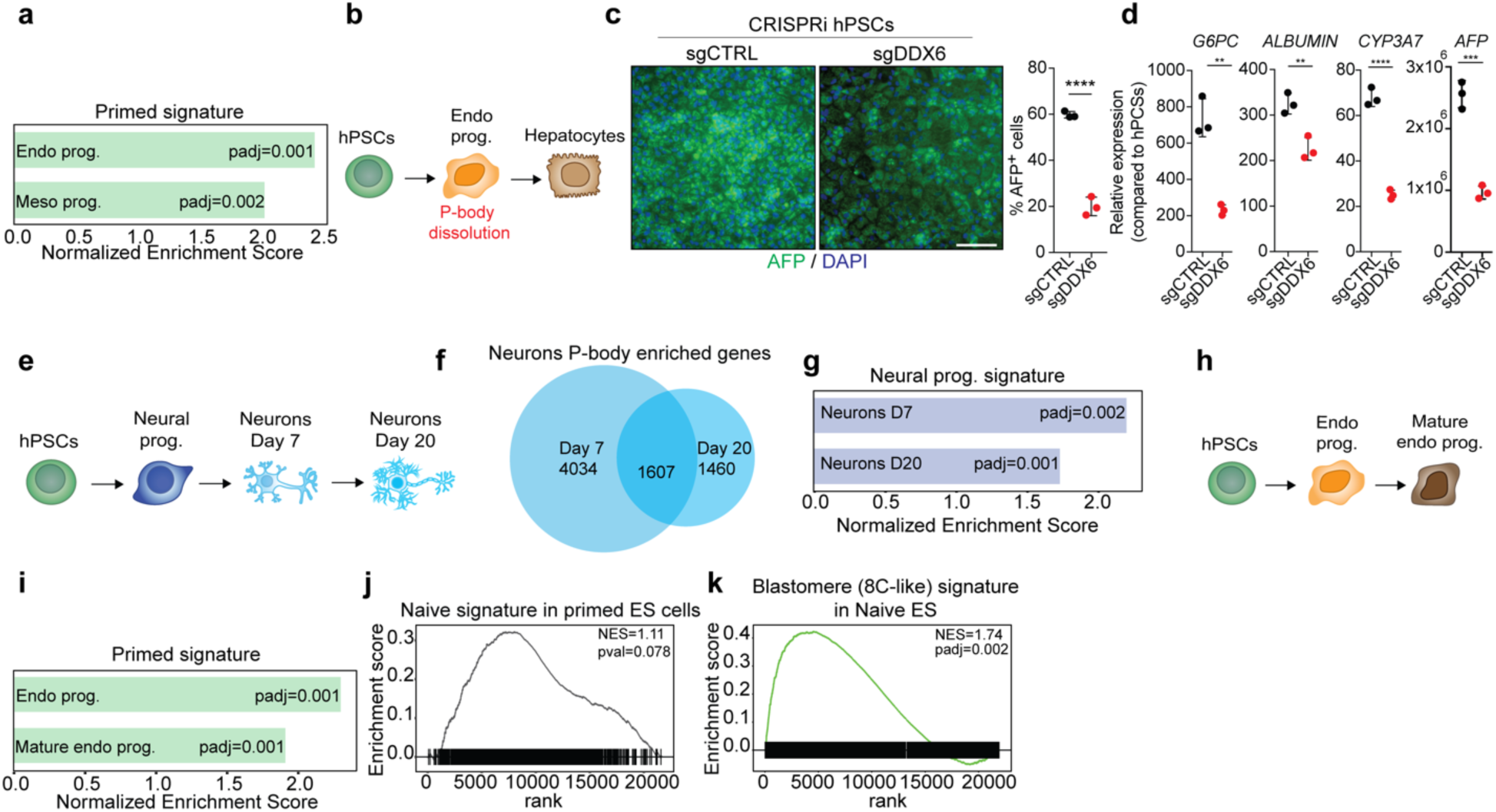
Transient sequestration of developmental stage-specific RNAs in P-bodies confers plasticity during early development. **(a)** Normalized Enrichment Score (NES) of primed human ES cell-related genes in purified P-bodies from endoderm and mesoderm progenitors. **(b)** A schematic of the strategy for P-body dissolution in human endoderm progenitors during their differentiation to hepatocytes. **(c)** Representative IF imaging of AFP (green) positive cells in hPSCs upon *DDX6* KO compared to control cells. Nuclei were counterstained with DAPI (blue) (scale: 100μm) (Left panel). Quantification of AFP^+^ cells upon *DDX6* suppression. Unpaired Student’s t-test, sgControl (n=3 fields), sgDDX6 (n=3 fields), mean ± s.d., ****: p<0.0001 (Right panel). **(d)** qRT-PCR analysis for the indicated genes in hepatocytes. Unpaired Student’s t-test, mean ± s.d. (*n=3*), (**p<0.01, ****p<0.0001). **(e)** A schematic of neuron maturation. **(f)** Venn diagram showing the overlap between P-body-associated mRNAs in neurons cultured for 7 and 20 days. **(g)** Normalized Enrichment Score (NES) of neural progenitor-related genes in neurons cultured for 7 and 20 days. **(h)** A schematic of endoderm progenitor maturation. **(i)** Normalized Enrichment Score (NES) of primed human ES cell-related genes in endoderm and mature endoderm progenitors. **(j)** GSEA analysis showing enrichment for a naïve signature in purified P-bodies from primed ES cells. **(k)** GSEA analysis showing enrichment for an 8C-like signature in purified P-bodies from naïve ES cells.

**Extended Data Figure 4.**
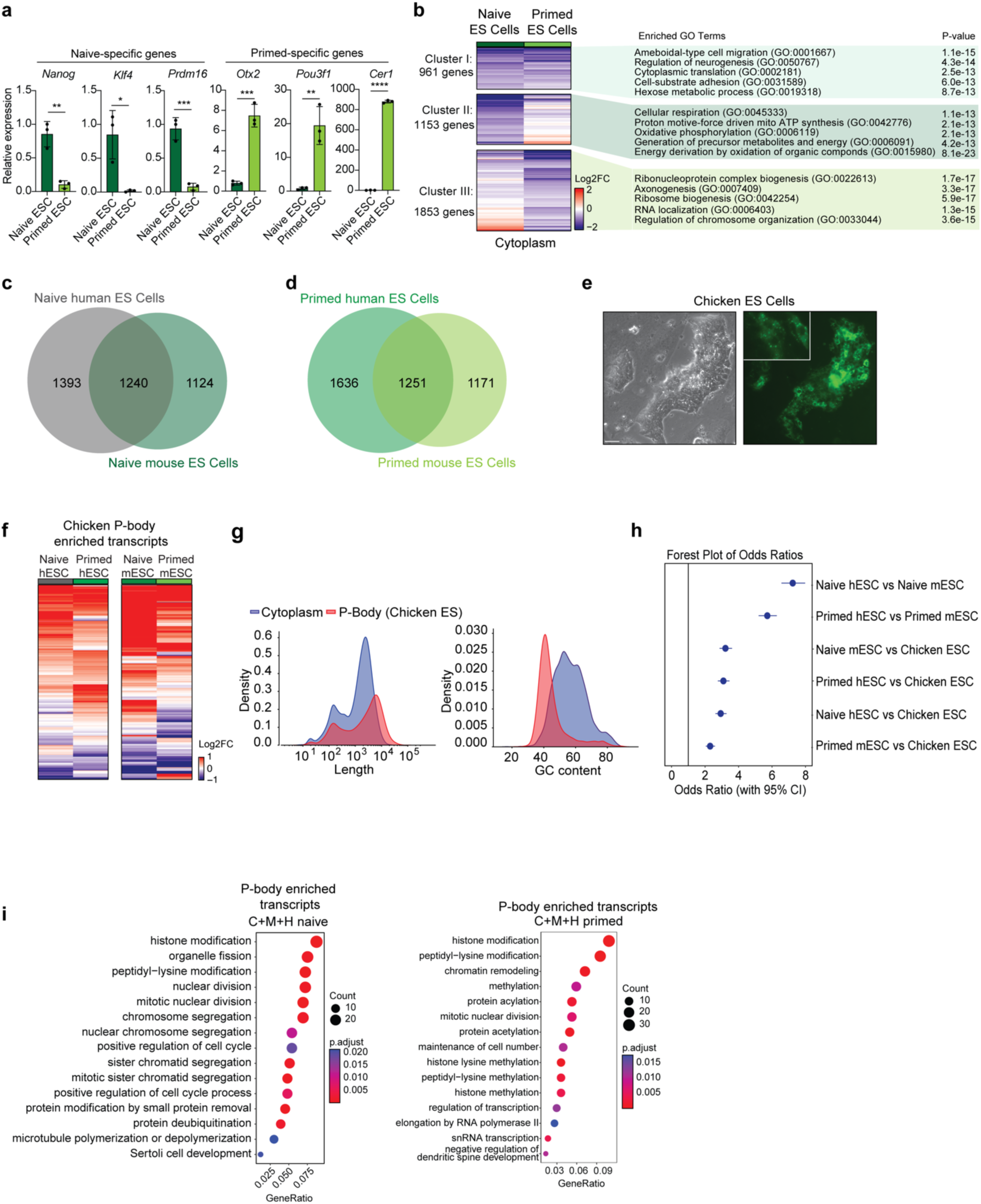
RNA sequestration in P-bodies is conserved across vertebrates. **(a)** qRT-PCR analysis of naïve-specific and primed-specific gene expression in naïve (n=3) and primed (n=3) mouse ES cells. Unpaired Student’s t test, mean ± s.d., *p<0.05, **p<0.01, ***p<0.001, ****p<0.0001. **(b)** Heatmap showing expression levels of differentially enriched mRNAs between cytoplasmic fractions of naïve and primed mouse ES cells, with GO pathway analysis of upregulated genes in each cluster. Gene number in each cluster is indicated in the figure (n=2, p < 0.05). **(c, d)** Venn diagrams showing the overlap between P-body enriched mRNAs in human and mouse naïve ES cells (c) and human and mouse primed ES cells (d). **(e)** Representative bright field image (left panel) and GFP expression (right panel) in GFP-LSM14A chicken ES cells (scale: 50μm). **(f)** Heatmap showing P-body enrichment in human and mouse primed and naïve cells for genes enriched in P-bodies from chicken ES cells. **(g)** Length (left panel) and GC content (right panel) density plots of mRNAs enriched in purified P-body and cytoplasmic fractions of chicken ES cells. **(h)** Forest Plot of odds ratios for P-body enriched genes in each cell type; error bars represent 95% confidence intervals (CI). **(i)** GO pathway analysis of P-body enriched mRNAs, showing common pathways between chicken ES cells and mouse and human naïve ES cells.

**Extended Data Figure 5.**
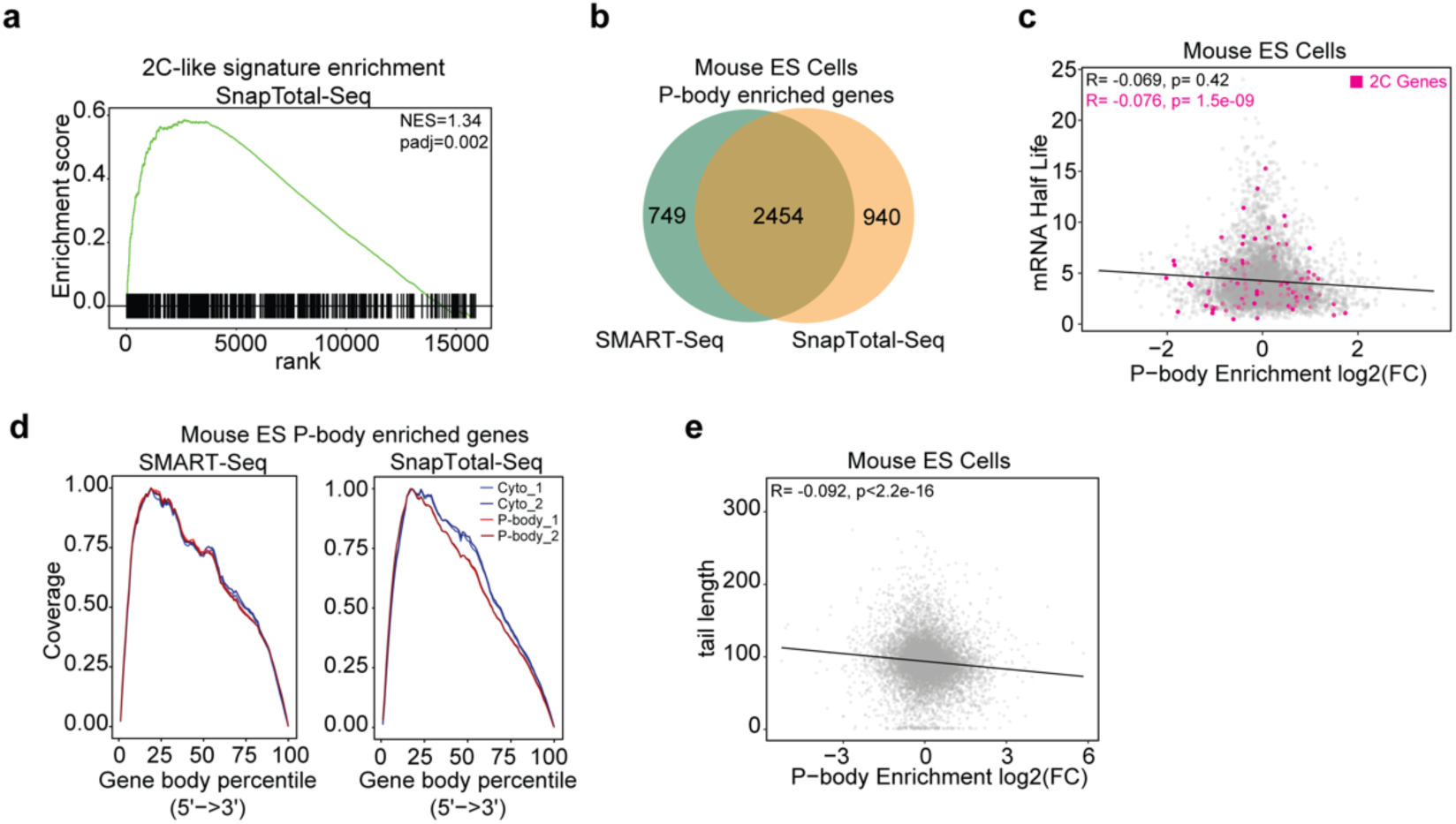
P-body-enriched RNAs in mouse ES cells are not preferentially degraded. **(a)** GSEA analysis showing enrichment for blastomere-related transcripts from mouse ES cells using SnapTotal-Seq. **(b)** Venn diagram showing the overlap between P-body-associated mRNAs in ES cells from SMART-Seq and SnapTotal-Seq. **(c)** mRNA half-life as determined in^90^ compared to P-body enrichment in primed mouse ES cells, 2C genes are highlighted in pink, Pearson correlation test. **(d)** Read coverage distribution over the gene body of the longest annotated isoforms of genes enriched in P-bodies in naïve mouse ES cells using SMART-Seq and SnapTotal-Seq. **(e)** Poly-A tail length as determined in^91^ compared to P-body enrichment based on SMART-Seq and SnapTotal-Seq, Pearson correlation test.

**Extended Data Figure 6.**
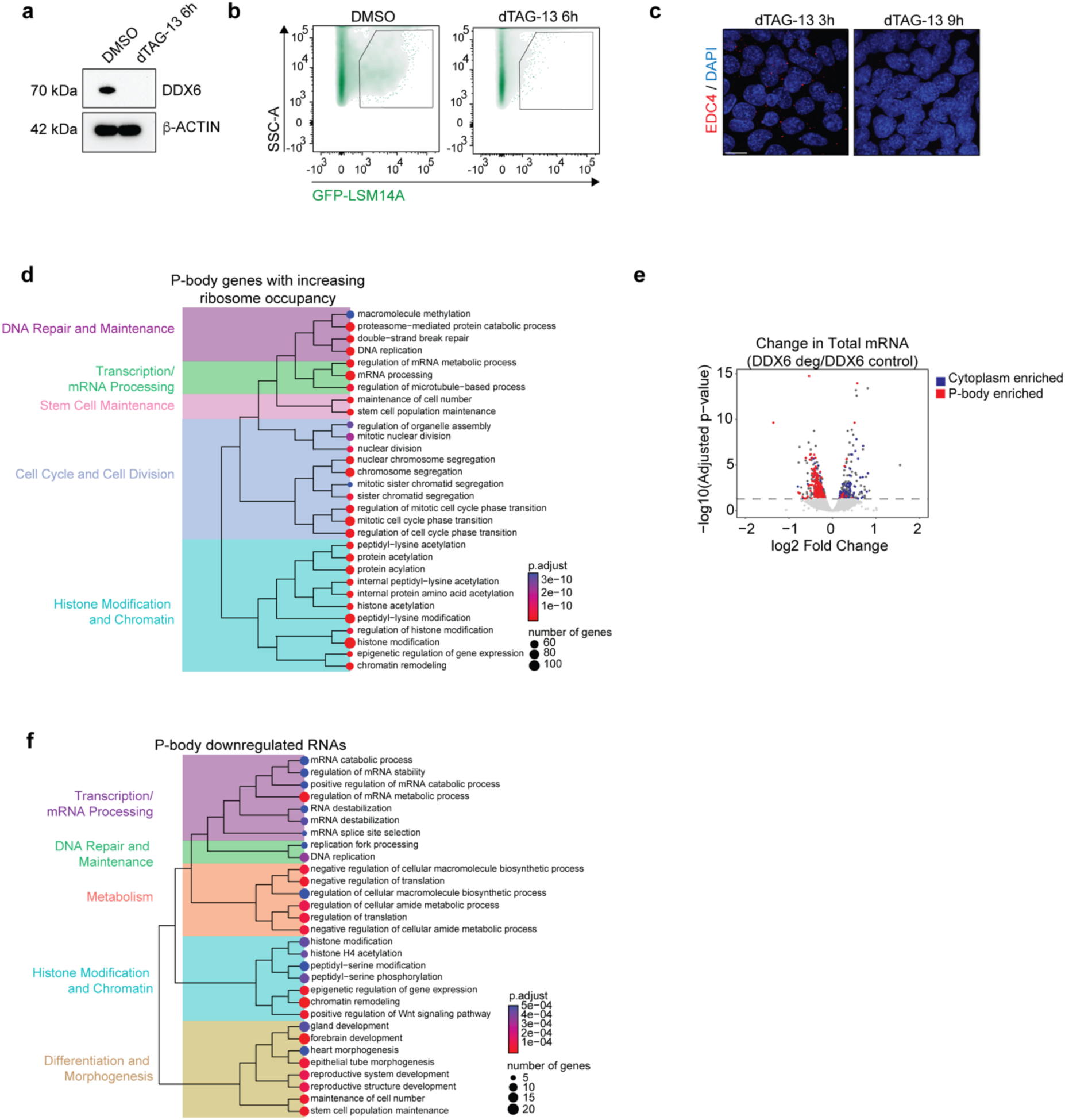
A degron system facilitates acute disruption of P-bodies. **(a)** Representative western blot showing DDX6 protein levels in *Ddx6*-FKBP12^F36V^ GFP-LSM14A mouse naïve ES cells, either untreated (DMSO) or treated with dTAG-13 for 6 hours. **(b)** Representative flow cytometry plots showing gating for GFP-LSM14A^+^ P-bodies in *Ddx6*-FKBP12^F36V^ GFP-LSM14A mouse naïve ES cells, either untreated (DMSO) or treated with dTAG-13 for 6 hours. **(c)** Representative IF imaging of EDC4 puncta (red) in *Ddx6*-FKBP12^F36V^ GFP-LSM14A mouse naïve ES cells, treated with dTAG-13 for 3 and 9 hours. Nuclei were counterstained with DAPI (blue) (scale: 10mm). **(d)** GO terms for P-body enriched genes with increased ribosome occupancy of *Ddx6-*FKBP12^F36V^ GFP-LSM14A mouse naïve ES cells following dTAG13 treatment for 6 hours). **(e)** Volcano plot of RNA-seq data depicting differential expression of total RNA fraction in *Ddx6-*FKBP12^F36V^ GFP-LSM14A mouse naïve ES cells following dTAG13 treatment for 6 hours, with P-body enriched genes in red and cytoplasm enriched genes in blue (n=3, p < 0.05). **(f)** GO terms of P-body enriched genes that showed downregulated gene expression in *Ddx6-*FKBP12^F36V^ GFP-LSM14A mouse naïve ES cells following dTAG13 treatment for 6 hours.

**Extended Data Figure 7.**
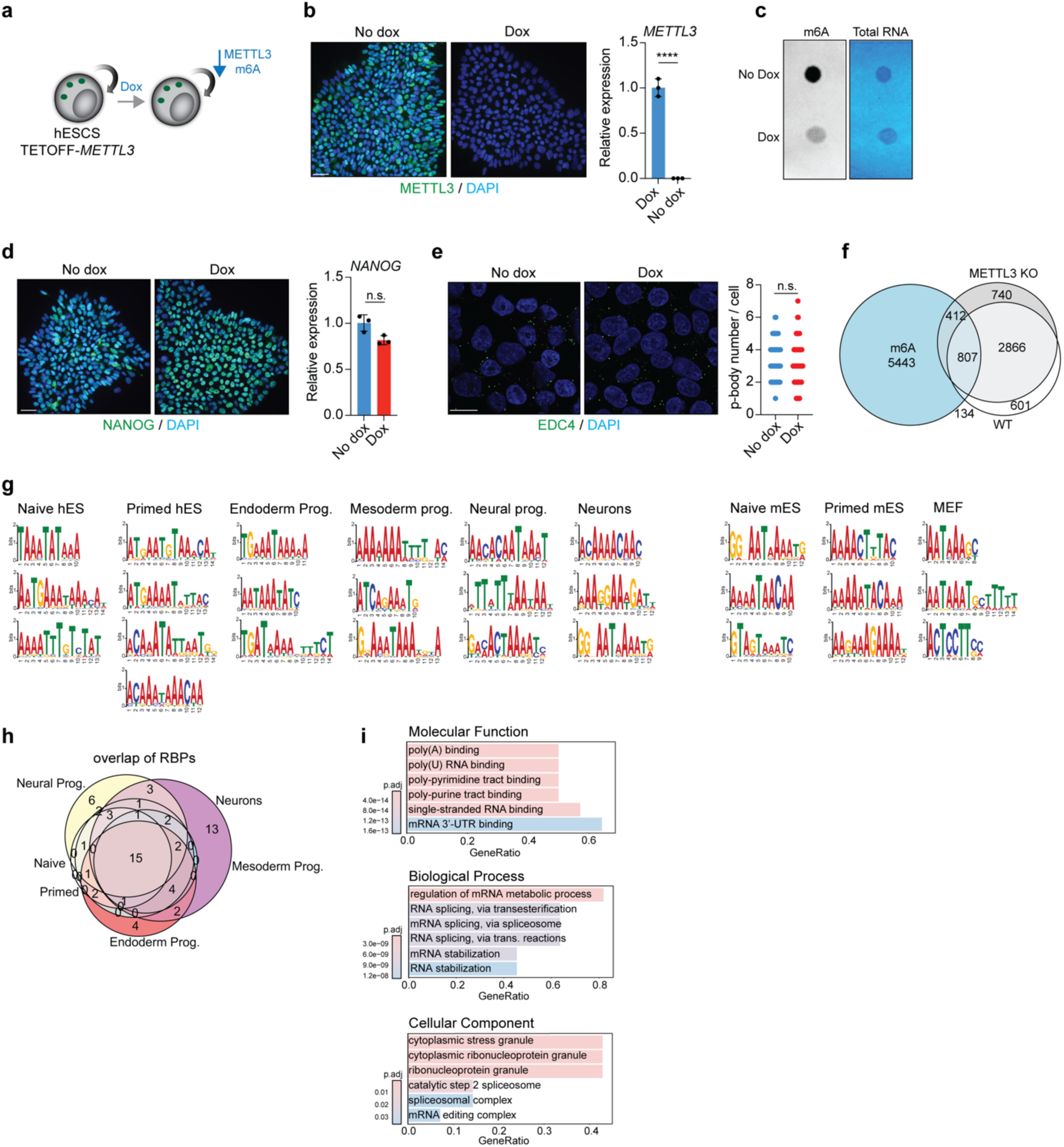
m6A RNA modification and RBPs do not drive cell-type specific transcript sequestration. **(a)** A schematic of the strategy for METTL3 suppression in TETOFF-*METTL3* human ES cells. **(b)** Representative IF imaging of METTL3 (green) in control (no dox) and METTL3 KO induced (dox) ES cells. Nuclei were counterstained with DAPI (blue) (scale: 50μm) (left panel). qRT-PCR analysis of *METTL3* expression in control (no dox) and METTL3 KO induced (dox) ES cells. Unpaired Student’s t-test, n=3, mean ± s.d., ****: p<0.0001 (right panel). **(c)** m6A RNA dot blot in in control (no dox) and METTL3 KO induced (dox) ES cells. Methylene Blue staining was used as loading control. **(d)** Representative IF imaging of NANOG (green) in control (no dox) and METTL3 KO induced (dox) ES cells. Nuclei were counterstained with DAPI (blue) (scale: 50μm) (left panel). qRT-PCR analysis of *NANOG* expression in control (no dox) and METTL3 KO induced (dox) ES cells. Unpaired Student’s t-test, n=3, mean ± s.d., ns: p>0.05. **(e)** Representative IF imaging of EDC4 puncta (green) in control (no dox) and METTL3 KO induced (dox) ES cells. Nuclei were counterstained with DAPI (blue) (scale: 10μm) (left panel). P-body number in control (no dox, n=56) and METTL3 KO induced (dox, n=56) ES cells (right panel). Unpaired Student’s t-test, mean ± s.d., ns: p>0.05. **(f)** Venn diagram showing the overlap between P-body-associated mRNAs in WT (no dox) and METTL3 KO induced (dox) ES cells, and m6A-methylated mRNAs in human ES cells. **(g)** Motifs enriched in the top 100 P-body enriched genes for each cell type, determined using STREME. **(h)** Euler diagram of the RNA Binding Proteins that were predicted to bind the motifs identified in (g) for each human cell type, determined by Tomtom with the Ray 2013 database. **(i)** GO terms for proteins that recognize the enriched motifs in all human cell types.

**Extended Data Figure 8.**
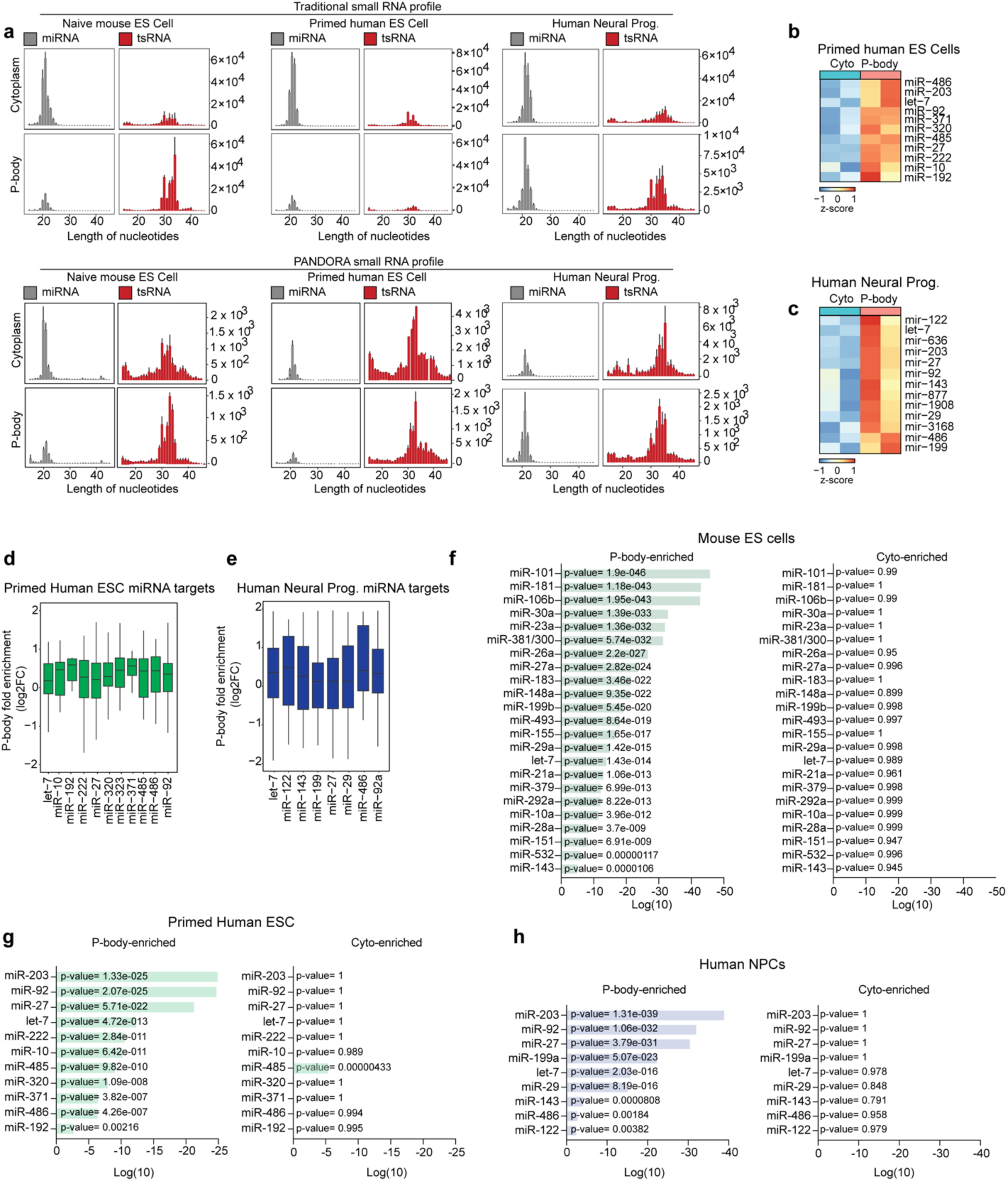
miRNAs direct selective sequestration of transcripts into P-bodies. **(a)** Changes in traditional (upper panel) and PANDORA (lower panel) small RNA distribution between purified P-body and cytoplasmic fractions in the indicated samples. **(b)** Heatmap showing expression levels of differentially enriched miRNAs between purified P-body and cytoplasmic fractions in primed human ES cells. (n=2, p<0.05). **(c)** Heatmap showing expression levels of differentially enriched miRNAs between purified P-body and cytoplasmic fractions in human neural progenitors. **(d)** Box plot showing miRNA-targets enriched in P-bodies of human primed ES cells (n=2, p<0.05). **(e)** Box plot showing miRNA-targets enriched in P-bodies of human neural progenitors. **(f-h)** MIENTURET predictions for miRNA targeting specifically P-body-enriched mRNA in mouse ES cells (f), primed human ES (g) and neural progenitors (h).

**Extended Data Figure 9.**
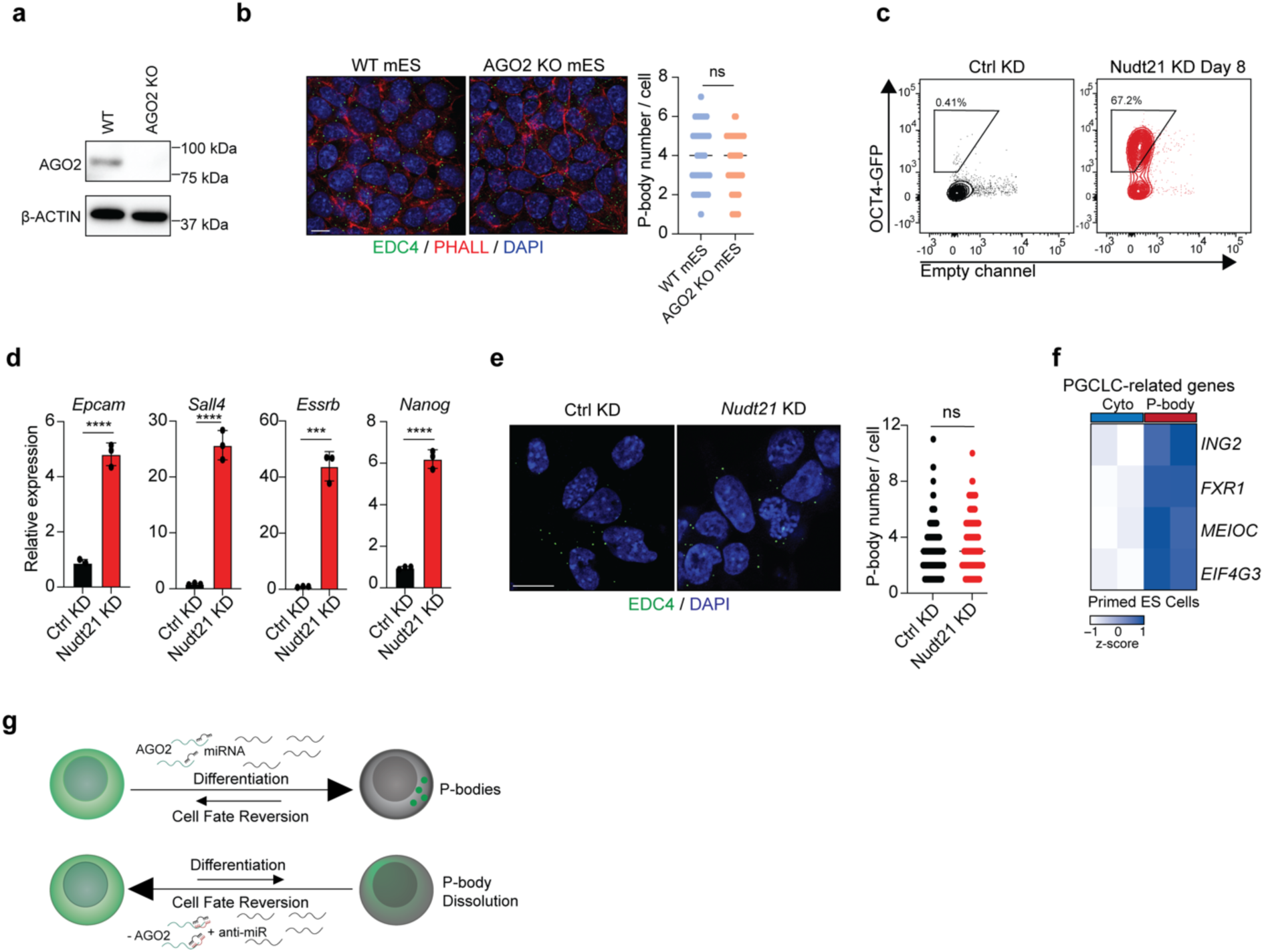
Uncoupling miRNA-mRNA interaction disrupts enrichment of transcripts into P-bodies. **(a)** Representative western blot showing AGO2 protein levels in WT and AGO2 KO mouse naïve cells. **(b)** Representative IF images of EDC4 puncta (green) in naïve and primed mouse ES cells. Cell membranes were labeled with Phalloidin (red), and nuclei were counterstained with DAPI (blue) (scale: 10μm) (left panel). P-body number in WT (n=60 cells) and AGO2 KO (n=60 cells) mouse naïve ES cells (right panel). Unpaired Student’s t-test, mean ± s.d., n.s.: p >0.05. **(c)** Flow cytometry quantification of OCT4-GFP^+^ cells in control and *Nudt21* KD reprogramming intermediates at day 8. **(d)** qRT-PCR analysis of the expression of ES-specific genes in control and *Nudt21* KD reprogramming samples. Unpaired Student’s t test, n=3, mean ± s.d., ***: p<0.001, ****: p<0.0001. **(e)** Representative IF imaging of EDC4 puncta (green) in control and *Nudt21* KD reprogramming samples. Nuclei were counterstained with DAPI (blue) (scale: 10μm) and P-body number in control (n=90 cells) and *Nudt21* KD induced (dox, n=90 cells) reprogramming samples (right panel). Unpaired Student’s t-test, mean ± s.d., ns: p>0.05. **(f)** Heatmap showing expression levels of differentially enriched PGCLC-related mRNAs between purified P-body and cytoplasmic fractions in human primed ES cells (n=2). **(g)** A model showing miRNA-mediated RNA sequestration in cell fate.

